# Logical modeling: Combining manual curation and automated parameterization to predict drug synergies

**DOI:** 10.1101/2021.06.28.450165

**Authors:** Åsmund Flobak, John Zobolas, Miguel Vazquez, Tonje S. Steigedal, Liv Thommesen, Asle Grislingås, Barbara Niederdorfer, Evelina Folkesson, Martin Kuiper

## Abstract

Treatment with drug combinations carries great promise for personalized therapy. We have previously shown that drug synergies targeting cancer can manually be identified based on a logical framework. We now demonstrate how automated adjustments of model topology and logic equations can greatly reduce the workload traditionally associated with logical model optimization. Our methodology allows the exploration of larger model ensembles that all obey a set of observations. We benchmark synergy predictions against a dataset of 153 targeted drug combinations. We show that well-performing manual models faithfully represent measured biomarker data and that their performance can be outmatched by automated parameterization using a genetic algorithm. The predictive performance of a curated model is strongly affected by simulated curation errors, while data-guided deletion of a small subset of edges can improve prediction quality. With correct topology we find some tolerance to simulated errors in the biomarker calibration data. With our framework we predict the synergy of joint inhibition of PI3K and TAK1, and further substantiate this prediction with observation in cancer cell cultures and in xenograft experiments.

## Introduction

Combining specific and targeted drugs in one therapy to fight disease increases chances of treatment success^1^. Drug combinations that together act in synergy are especially attractive because they allow for pushing treatment effects beyond those obtainable by each drug alone^2^, with drug dosages that can be well below levels where individual drugs begin to cause adverse effects. In addition, synergistic drug combinations may have reduced side-effects by improved selectivity in a specific biological context, for instance by allowing targeting of only certain cell types in an organism^3^. Lastly, searching for new combination therapies has the additional benefit that already approved drugs can act beneficially in novel combinations, and thus even allow bypassing initial drug development phases.

While the development of rational drug combination treatment has become a major priority due to hopes of increased treatment potency, a grand challenge remains in dealing with their identification in the vast potential drug combination space. Currently, more than two hundred drugs have been approved by the FDA to treat cancer (https://www.cancer.gov/about-cancer/treatment/drugs). The testing of drugs in all combinations with other drugs in a panel needs assays in numbers that increase exponentially with increasing drug numbers. Even a modestly sized drug panel of 150 drugs corresponds to over 10.000 pairwise drug combinations. Testing high numbers of drug combinations in high throughput screens on cell lines or other patient-specific model systems is costly and at some point prohibitively expensive and cumbersome. Therefore, help is needed from *in silico* drug effect simulations to produce high quality predictions that can guide drug combination screens or therapy choices for testing in cell lines or patients. *In silico* simulations may help identify those combinations that are unlikely to produce synergies, which can be of significant help to reduce the large experimental search space that otherwise would need to be covered in exhaustive screens. As many drug synergies can be seen as emergent properties arising from molecular causality networks, analytical frameworks from computational systems biology seem to be well suited to the task.

Several mathematical frameworks have already been tested to mechanistically model drug combination effects, including continuous, discrete, and hybrid modeling approaches. Published approaches generally depend on molecular causalities downloaded from prior knowledge databases, extracted from large-scale data, or obtained by a combination of the two. Based on a dataset capturing proteomic responses of 14 targeted drugs, Miller et al. used ordinary differential equations to study mechanisms of synergy between inhibitors of CDK4 and IGF1R, revealing that the mechanisms rely on the activity of AKT^4^. Nelander et al. explored ODE models derived from observations on phospho-proteins and cell cycle markers following 21 pairwise applications of targeted drugs, with the aim to use best-performing pairs for design of new combination therapies^5^. In a semi-qualitative modeling approach, Klinger et al. used a perturbation dataset for MAPK, PI3K and NF-κB signaling to inform a model showing that combined inactivation of MEK and EGFR could inactivate endpoints of RAS, ERK and AKT signaling^6^. Jin et al. explored enhanced Petri nets to describe molecular processes for the synergy of an EGFR inhibitor (gefitinib) with chemotherapy (docetaxel) and identified KRT8 as a candidate gene to explain the synergy^7^. However, all of these approaches rely on extensive and costly combinatorial drug perturbation data, be it transcriptomic, proteomic, viability etc., for describing mechanisms of synergy, and therefore they require vast investments in data production and do not provide a feasible solution for the testing of the large drug combination space.

In order to reduce dependence on *a priori* perturbation experiments, attempts have been made to predict drug synergies from data obtained in a marginal experimental search space, rather than the full combinatorial space. Fröhlich et al. used ODE models informed by transcriptomic and viability data to predict drug combination responses, finding that highly accurate predictions could be produced for those drugs for which they had viability response data^8^. In the DREAM7 - NCI-DREAM, Drug Sensitivity and Drug Synergy Challenges^9,10,11^ (NCI-DREAM), pairwise drug responses were predicted from response data obtained for each drug alone. The best performing teams in the NCI-DREAM challenge obtained a probabilistic concordance (PC) index of 0.61, on a scale ranging from 0.9 (perfect prediction) to 0.1 (perfect opposite prediction). Although this is better than random (PC index of 0.5), it clearly illustrates that obtaining accurate synergy predictions is far from trivial, due to a variety of reasons that will be discussed in this paper. In the more recent AstraZeneca-Sanger Drug Combination Prediction DREAM Challenge^12^ (AZS-DREAM), one of the aims was to develop and demonstrate drug combination response predictability independent of extensive perturbation data. The development of such powerful prediction approaches has the advantage of being relevant not only to preclinical drug screens, but also to bed-side applications. Drug perturbation data clearly will not be trivial to obtain for individual patients, unless patient-derived experimental assays that mimic patient responses can be developed (e.g. xenografts, explants etc.). Sobering results from the AZS-DREAM challenge showed that most teams had balanced accuracies of 0.5-0.6, with the best performing team obtaining a balanced accuracy of only 0.69.

With the availability of training data, machine learning algorithms have also been explored to predict drug synergies^13, 14, 15^. However, major limitations of such approaches are the lack of mechanistic insight^16^, and dependence on high quantities of training data. Despite some increase in their availability, large scale datasets on drug responses for machine learning to predict combination effects are still largely missing, in part due to great experimental complexity and high economic cost. Whereas efficient *in silico* therapy based on patient-specific models ultimately should be integrated into the clinical decision process, here we further investigate the performance of logical modeling of drug response of cancer cell lines. In order to reduce the dependency on large training datasets, we explore the use of cell lines measurements of biomolecules obtained at a single time point at steady state proliferation. To reduce data dependency and to improve mechanistic insights, these measurements are combined with prior knowledge to construct logical model ensembles to simulate drug combination effects. Logical model building is known to require meticulous involvement of curators and bioinformaticians, with substantial commitment to manual tinkering of models before the behavior of one model matches that of its experimental counterpart. We have previously published logical models of cancer cell lines, named CASCADE 1.0, CASCADE 2.0 and CASCADE 3.0, which demonstrate the potential of logical modeling for the prediction of drug synergies^17, 18, 19^. However, the curation effort required to assemble a cancer signaling network and the dedicated interactive efforts needed to optimally modify logical rule definitions becomes a clear obstacle when constructing larger models.

If patient specific logical models are to be used routinely, such logical models should be trivial to construct for any cell line or patient-derived cell culture, and for any repertoire of targeted drugs. We therefore set out to automate processes required to calibrate a logical model from a set of molecular causative statements, i.e. a prior knowledge network. A software pipeline developed to that end would have to 1) assemble a network topology from structured data obtained from prior knowledge databases, 2) interpret baseline cancer cell line biomarker data into a signaling entity activity score, 3) calibrate generic logical models, created from prior knowledge data, by modifying logic equations to match the observed activity scores, and 4) predict phenotypic consequences of combinatorial interventions to the simulated model behavior. Our software solution for realizing points 3 and 4 is available at https://github.com/druglogics. We use a genetic algorithm to automatically parameterize a set of logic equations representing cancer growth-promoting signaling in the AGS gastric adenocarcinoma cell line. We demonstrate our approach by reproducing results from a previous manual effort and next test its utility with a larger model that was benchmarked against a dataset from a drug effect screen of 153 drug combinations. Experiments that simulate different levels of curation quality and biomarker data quality indicate the need for a reliable PKN, while still allowing for model improvement by network link pruning and parameter optimization.

## Methods

### Logical modeling

Logical models rely on the formalism initially proposed by Stuart Kauffman^20^ and René Thomas^21^. Their approach first defines a regulatory graph consisting of nodes representing signaling entities (model components), and signed and directed edges representing regulatory interactions that connect signaling entities. The activities of all model components are then associated with the Boolean values ‘True’ and ‘False’, represented by 1 and 0, corresponding to activity and inactivity, respectively. This dichotomy of activity levels can be interpreted as activity being above or below a “threshold”: a component is “active” when its activity level is sufficiently high to influence a target component’s activity levels. In model simulations, specific model components can be defined as ‘output nodes’, whose activity values serve as a proxy for a phenotype of interest. This allows us to compute an overall ‘growth’ value in our simulations, by integrating all activity values of model output nodes, and scaling this sum to [0..1]. For example, if the anti-survival model output nodes CASP8, CASP9 and FOXO3 are inactive and the pro-survival model output nodes MYC, CCND1 and TCF7 are active, the global output ‘growth’ is 1.0. Inhibitory effects of a drug are simulated by fixing the drug target to an activity level of 0, i.e. simulating a block in signaling activity of the drug target node.

Model attractors were determined using the software bioLQM, which provides a fast algorithm for finding stable state(s) and complex attractors^22^. For some of the simulations that were too computationally taxing, we used a modified version of the algorithm BNReduction^23^, which allows the identification of single stable state phenotypes that are most prevalent with our self-contained CASCADE topologies.

#### Model calibration by parameterization optimization

A genetic algorithm is applied to automate model parameterization, as follows:

The input for model calibration consists of:

- Interactions: binary signed and directed interactions (SIF format^24^).
- Steady state: Boolean vector containing states of nodes in input interactions, where nodes that should be active are assigned the value 1, and nodes that should be inactive are assigned the value 0. For nodes whose state cannot be determined a dash (−) can optionally be used to explicitly declare that the node state is undetermined. This steady state vector will be used for evaluating the correctness of a model’s stable state.

The output from an automated model calibration is an ensemble of models with a stable state optimized to match the input steady state.

To run the parameter configuration, interactions are first assembled to logic equations, based on a default equation^25^ relating a node with its regulators, for instance:

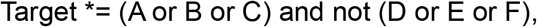

where activating regulators A, B and C of a target are combined with logical ‘or’ operators, and inhibitory regulators D, E and F are combined with ‘and not’ operators to determine the state of the target node in the next time step. The operator ‘and not’, which directs the integration of activating and inactivating regulators, is referred to as the link operator in this manuscript. The topology is ‘self-contained’, meaning that any regulator is itself regulated by one or more components from within the network topology, effectively meaning there are no ‘user-controlled’ inputs to the network through e.g. hormone receptors.

Next, a genetic algorithm is used to iteratively refine the parameterization to produce a logical model with a stable state matching the specified input steady state. First, an initial generation of models is formulated, where a large number of mutations to the parameterization is introduced: randomly selected equations are mutated from “and not” to “or not”, or vice versa. For each model a fitness score is computed: each matching Boolean value between the vector of a stable state and the steady state improves the fitness score. Models without a stable state have a fitness of zero. Models with *n* stable states obtain a final fitness after integrating all Boolean values in a stable state vector and dividing the resulting overall fitness by *n*, thus penalizing models with multiple stable states.

For each generation, a user-defined number of models were selected for populating the next generation of models. For our simulations three models were selected, specifically the ones that achieved the highest fitness scores in each generation, to populate a next generation of 20 models. First, crossover was performed, where each selected model would exchange logic equations with other selected models (including itself, thus also enabling asexual reproduction). Then a number of mutations were introduced as described above. For our simulations up to three mutations were introduced. Before a stable state was obtained, the number of mutations introduced per generation was increased by a user-defined factor. The large number of mutations in the initial phase ensured that a large variation in parameterization could be explored. For our simulations we chose the factor to be 1000, effectively ensuring that the initial generations were randomly sampled from all possible model configurations. Evolution was halted when a user-specified threshold fitness was reached. In case this fitness could not be reached, evolution was halted when a user-defined maximum number of generations had been spanned. For our simulations we allowed for a maximum of 20 generations.

#### Model calibration by topology optimization

In order to introduce variations to the topology, the genetic algorithm modified a whitelist and a blacklist of regulators of the prior knowledge network, while always preserving at least one regulator for each target, so as to not break the self-contained property of the network. Based on the same formula as shown above,

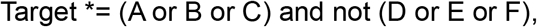

this means that the genetic algorithm takes out some subset of the regulators A, B, C, D, E or F (blacklisting). After a regulator has been eliminated, the genetic algorithm is also allowed to bring back regulators originally defined in the PKN (whitelisting). For our simulations, during the initialization phase we introduced 50 such topology mutations and when models with stable states were found, we reduced this number to 10, so as to not severely reduce the PKN edges.

#### Model simulation and synergy prediction

After repeating evolution a specified number of times, model ensembles were analyzed in a third step of the software pipeline, as follows:

The input to model simulation and synergy prediction consists of:

- An ensemble of logical models
- A drug panel: List of drugs and their target node(s) in the model
- Perturbations: the perturbations to be analyzed.
- Model output nodes with weighted score to evaluate global output (i.e. ‘growth’)

Output from model simulation and synergy prediction:

- Drug synergy predictions from ensembles of models

For each model, all perturbations specified were simulated. For each perturbation, the drug panel was consulted to fix the state of the specified node(s) to the value 0 (node state could in principle also be fixed to the value 1 for a drug that *activates* a signaling entity, but this feature was not used here as all drugs *inhibit* nodes in the model, thereby representing inhibition of their target in a cell). After simulating a perturbation, the global output parameter ‘growth’ was computed by integrating the weighted score depending on the states of model output nodes. For example, if two output nodes A (weight 1) and B (weight −1) were found to have the states A=1, B=1 for a perturbation, the global output would evaluate to A_state_*×*A_weight_ + B_state_*×*B_weight_ = 1*×*1 + 1*×*(−1) = 0. This value was then scaled from 0 to 1 based on the theoretical minimum and theoretical maximum ‘growth’, for this example, the range [-1,1], the global output would be 0.5. The scaled global output (‘growth’) was then used to compute synergies (see below).

All steps of the software pipeline were implemented in the OpenJDK Java v1.8 language and run on Linux 4.15.0-122-generic/Ubuntu 18.04.4 LTS. The pipeline can be accessed at https://github.com/druglogics. For an extensive documentation of the methods used in this work, see https://github.com/druglogics/ags-paper.

#### In silico definition of synergy

Synergy is defined as an additional response beyond what is expected from a reference model of drug combination responses. Both for *in silico* simulations and *in vitro* experiments an observed combination effect can be formally defined as the effect *E* observed for two drugs *a* and *b,* where *E(a,b)* is the observed effect in a combination experiment, *A(a,b)* is the drug combination effect expected from each individual drug’s properties as based on a reference model for combination responses, and *S*(*a*,*b*) is any difference between the observed and the expected drug combination effect, such that *E(a,b) = A(a,b) + S(a,b)^26^*. In the case of excess effects observed for a combination, *S*(*a*,*b*) is positive and synergy is called, and conversely for attenuated effects, *S*(*a*,*b*) is negative and antagonism is called. Finally, for drug combinations where *E*(*a*,*b*) equals *A*(*a*,*b*), the drug combination effect can fully be anticipated by each drug response independently, and neither synergy nor antagonism is called.

In model simulations the expected drug combination response is defined as the product of the two global output ‘growth’ values for each single drug, similarly to the Bliss independence^27^ synergy metric used in lab experiments: when a combinatorial perturbation in simulations is found to predict a lower growth than expected, i.e. *growth(a,b) < growth(a)* * *growth(b),* the combinatorial perturbation response is declared synergistic.

### Gold standard synergies

Our previously published dataset of targeted drug combinations^[17]^ was used to benchmark the algorithms. The drugs included comprised the inhibitors 5Z-7-oxozeaenol (5Z), AKTi-1,2 (AK), BIRB0796 (BI), CT99021 (CT), PD0325901 (PD), PI103 (PI), PKF118-310 (PK), JNK Inhibitor XVI (JN), BI-D1870 (D1), BI605906 (BIX02514) (60), Ruxolitinib (INCB18424) (SB), SB-505124 (RU), D4476 (D4), KU-55933 (KU), 10058-F4 (F4), Stattic (ST), GSK2334470 (G2), GSK-429286 (G4), P 505-15 (P5). For the drug synergy calling in the 153 combinations drug screen, three curators were asked to evaluate growth curves and decide on which showed interesting combination effects that could have warranted further investigations. A consensus list was then used to identify a threshold for drug synergy assessment using the software CImbinator^28^ and configured to compute synergies per the Bliss metric. The analysis identified six drug synergies (AK-BI, PI-D1, BI-D1, PI-G2, PD-PI, 5Z-PI). Note that two drug synergies in the drug screen performed in 2015 were not captured by this analysis, probably relating to the different readouts used in the drug screen performed in 2019 (xCELLigence and CellTiter Glo, respectively).

### Normalization

Normalization of synergy predictions was performed by computing the exponential fold change for the ratio of output from models *calibrated* to steady state biomarker data (x) to output from models calibrated to a *random* yet proliferative phenotype (y):

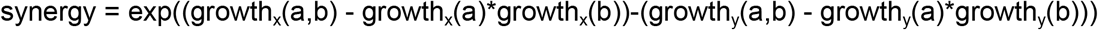

Our random proliferative phenotype corresponds to a cell with all anti-apoptotic signals inactivated, and at least one prosurvival signal active.

### Mouse xenograft experiments

40 female Balb/c mice 4-5 weeks old (Taconic) were inoculated with two million AGS cells subcutaneously in the right dorsal flank. Cells were mixed with Matrigel to improve probability of successfully establishing a xenograft model: in a small pilot (n=3) we observed that none of three mice injected with cells in medium (DMEM) developed tumors, while two of three mice injected with cells in medium and Matrigel developed tumors. 100 μl of cell suspension in HAM’S F12 medium (Invitrogen, Carlsbad, CA) with 10% fetal calf serum (FCS; Euroclone, Devon, UK), and 10 U/ml penicillin-streptomycin (Invitrogen) was mixed with 100 μl of ECM Gel from Engelbreth-Holm-Swarm murine sarcoma (Sigma-Aldrich). After four weeks, minuscule but palpable tumors had formed in 30 mice, which were randomized to four groups and subjected to treatment: 1) 5Z-7-oxozeaenol (3 mg/kg/d), 2) PI103 (5 mg/kg/d), 3) 5Z-7-oxozeaenol (3 mg/kg/d) + PI103 (5 mg/kg/d), 4) vehicle. Randomization ensured that average tumor volume was similar in the four groups. Weights of mice ranged from 14.9 to 20.0 grams at onset of treatment, with average weight 17.66 g and standard deviation of 1.06 g. All mice received the same dose of drugs, and the dose was adjusted for a body weight one standard deviation below average, i.e. 16.6 g. Drugs were diluted in medium with 40% DMSO for a total injection volume of 250 μl and injected intraperitoneally three times per week for a total of seven injections. Maximum (a) and minimum (b) tumor diameters were measured twice weekly with a caliper, and the volume V of the tumor was estimated from the formula V = 0.5 a × b^2^.

## Results

For reliable simulation of drug responses of cancer cell lines, computational models must adequately represent the regulatory network (topology) underlying cell fate decisions, meaning that high quality molecular causal relationship data must exist and be converted to regulatory graphs. In addition, the activity states of molecular regulatory components must be measured, demanding high quality biomarker data. From the regulatory graph the response of components to upstream source nodes and influence on downstream target nodes needs to be specified in the form of logical rules and calibrated so as to accurately represent the biological decision mechanisms of these cells in a Boolean framework. Finally, good benchmarking data must exist to evaluate the performance of the model, e.g. for our purposes in the form of drug synergy data, see figure 1A.

**Figure 1A:**
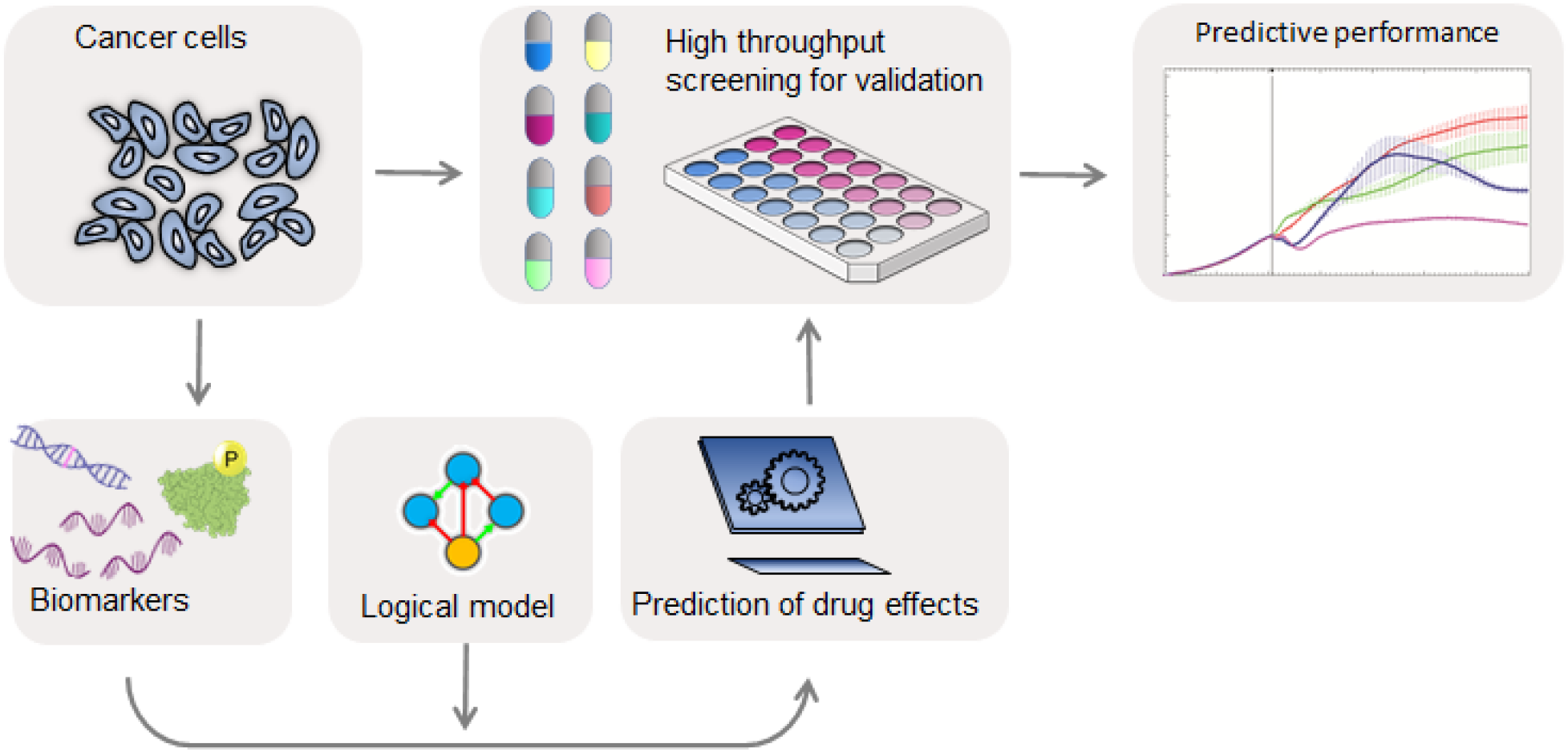
Overview of the drug synergy prediction platform. Cancer cells are analyzed for biomarkers used to define logical models that can be used to predict drug synergies. Model predictions are tested by benchmarking against high throughput drug screens.

### Design of an automated model parameterization module

Previously we have shown the feasibility of logical model predictions of drug synergies^17^ using the cancer cell line AGS, chosen due to known deregulations of several core cell survival signaling pathways. A Prior Knowledge Network (PKN) was curated to represent these signaling pathways, and converted to a set of mathematical rules formulated in Boolean logic. This model, available from https://github.com/druglogics/cascade as CASCADE 1.0 (Fig. 2A), consists of 75 nodes representing cancer signaling entities and 149 edges representing regulatory interactions, and it could predict five synergies of which four were experimentally confirmed^17^. Since our model is based on prior knowledge, amenable to interpretation by molecular biologists, the model also can be used for inspection as to which signaling pathways are important for particular drug response observations, e.g. we have suggested that FOXO signaling was crucial to the drug synergy effect of joint PI3K and MEK inhibition^17^. Whereas for many of the logical rules the definition of the logical operators (AND, OR, NOT) was more or less evident from literature and database knowledge, analysis of Boolean model attractors indicated that some logical rules needed further manual optimization in a stepwise manner so that ultimately the model stable state behavior matched the observed pattern of signaling entities at steady state in proliferating AGS cells. We now report on how we automated and generalized these steps required to parameterize an *in silico* model of a cancer cell line, by employing a genetic algorithm for deriving logical rules from prior knowledge and steady state signaling observations, see figure 1B. From a curated network topology a set of standardized logic equations are obtained by defining one logic equation for each model target node, with model source nodes as operands^25^. For example, if protein T is activated by proteins A and B, while protein C inhibits protein T, the equation could read as T = (A *OR* B) *AND NOT* C. Subsequently, the parameterization (choices of logical operators) is optimized by a genetic algorithm, specifically modifying the AND NOT/OR NOT parameter. The genetic algorithm iteratively modifies the parameterization of a small subset of the equations, and selects best performing models for defining a new generation of candidate models. Best performing models are chosen based on their compliance with reproducing known baseline cell signaling states, as far as the available cell line data allows it. Evaluating fitness from a match with baseline observations also means that our models are defined independently of perturbation data. Our software solution for automatic parameterization is available at http://github.com/druglogics/.

**Figure 1B:**
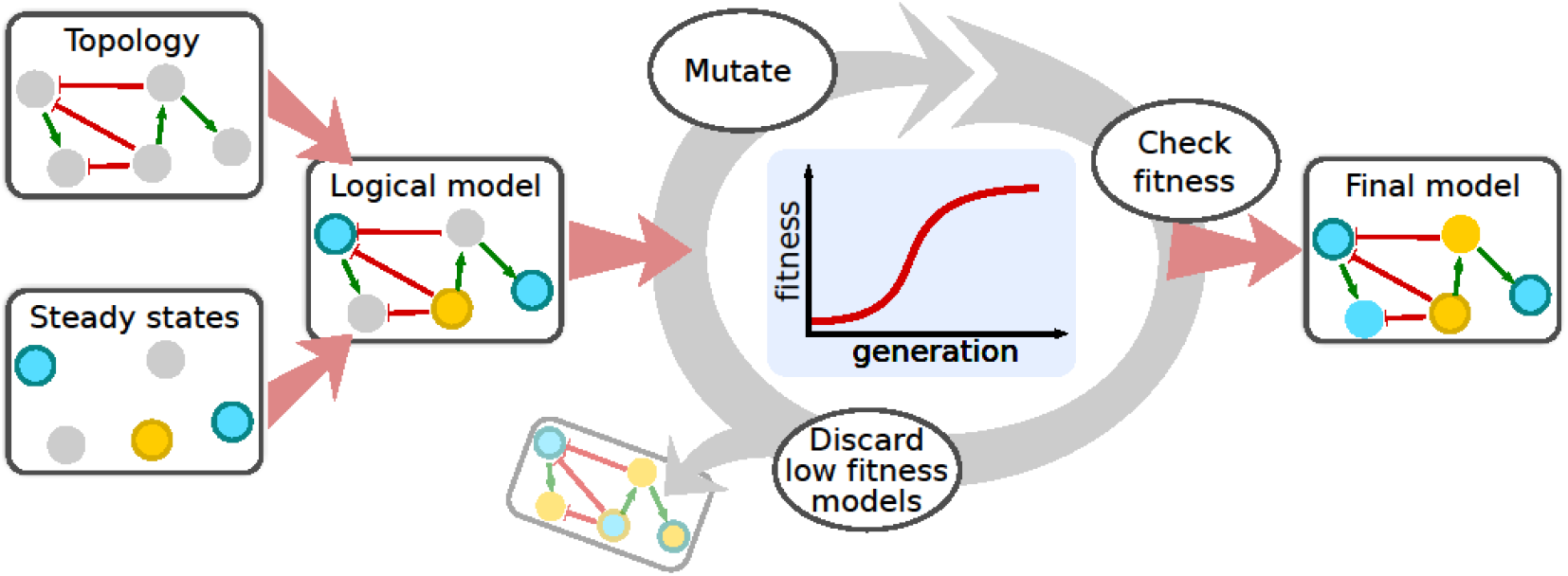
A genetic algorithm optimizes logical models to cancer cells. A prior knowledge signaling topology representing molecular causal interactions is taken as input to define logical models with predefined rules as initial logic equations. A genetic algorithm will iteratively randomly choose logical rules by mutating the AND/OR configuration, thereby re-parameterizing a logical model until a model shows a maximum compliance with steady state signaling observations (biomarkers). This procedure is performed in parallel for hundreds of models until an optimized ensemble of models is available for drug synergy prediction.

**Figure 2A:**
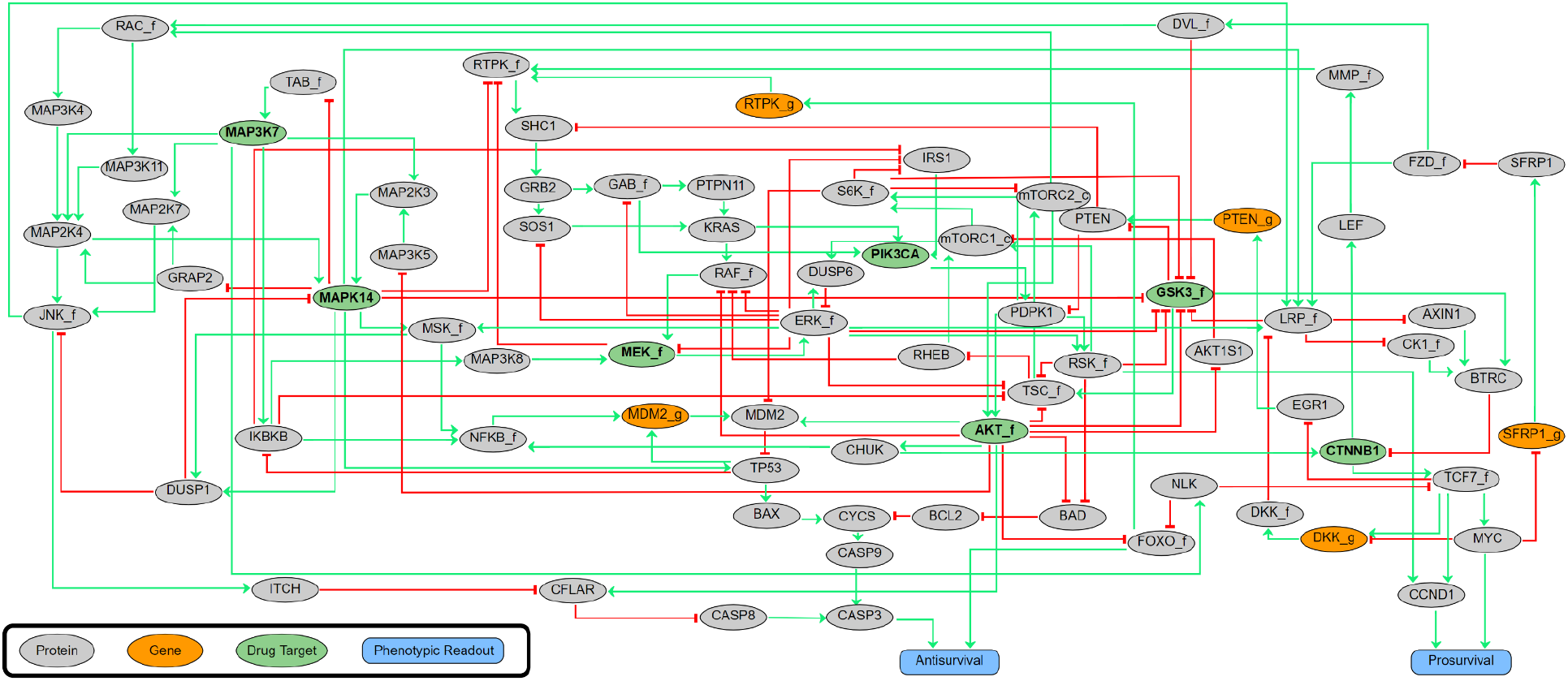
The CASCADE 1.0 prior knowledge network. 75 signaling nodes with signed and directed regulatory influences annotated (activating interactions in green, inhibiting interactions in red). All signaling components receive input from other signaling components from within the network, and ultimately influence the two phenotypic output nodes Antisurvival and Prosurvival.

### Automated logical rule definitions perform on par with manually curated rules

To compare results from our manually constructed logical rules with automated rule definitions, our software translates the graph, as encoded in a SIF file format, to a set of 75 logic equations in a standardized format^25^. Logic equations are then optimized using a genetic algorithm in a process where the fitness of each model is calculated by comparing the matches of its stable state nodes with observed activities of signaling entities for proliferating cancer cells. For the AGS cells, this process comprised 20 generations, in which each generation received mutations to a small subset of logic equations iteratively, with 20 models per generation tested for fitness. In order to adequately cover the space of local optima, the evolutionary process was repeated 50 times and the three best performing models from each evolution were retained, which resulted in an ensemble of 150 models. A theoretical maximal fitness of one would be reached if all nodes have a state matching the observed activity state of the corresponding protein. As can be seen in figure 2B, the population average fitness of each generation increases exponentially before plateauing at a fitness close to the theoretical maximal fitness, per Holland’s Schema Theorem^*29*^, indicating that the theoretical models can be parameterized so as to be compliant with experimentally observed signaling states. While a genetic algorithm cannot guarantee a global optimum, our results clearly indicate that we quickly achieve convergence to a local optimum.

**Figure 2B:**
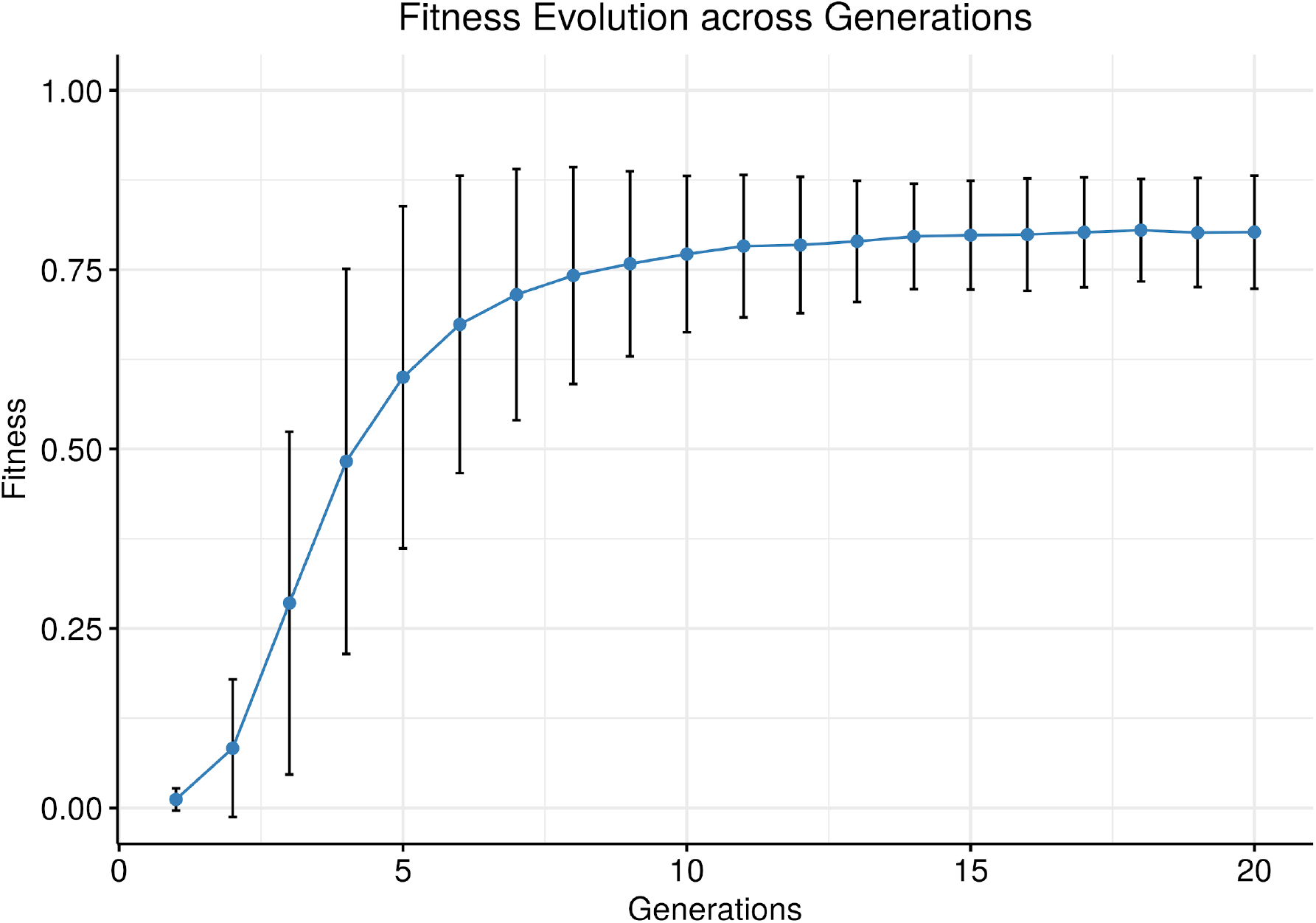
Evolution of fitness of calibrated models. Overall fitness is plotted as a function of generation, with average fitness and standard deviation indicated. The data for this figure was produced by running the genetic algorithm for 1000 simulations, with 20 generations per simulation and 20 models per generation. We observe that the average fitness and standard deviation follow a sigmoidal increase and stabilize after 10-15 generations. The persistence of the standard deviation across generations including those late in the evolution shows that new models still explore variations to the model parametrization while selection keeps the fitness score of the trained models at a constant plateau.

Whereas these model ensembles provide the testing ground for the *in silico* drug effect simulations, it is to be expected that certain motifs of the network topology itself will create ‘blind spots’ resulting in some synergies to be impossible to predict. For example, if two directly sequential signaling nodes are targeted by two different drugs, while no other influences from other signaling entities are allowed by the topology, then these two drugs cannot be predicted to act synergistically in our logical modeling framework. To remedy such situations extensions to the prior knowledge network are necessary, or conversion to non-discrete modeling. On the other hand, if two drug targets are active and are the only (activating) source nodes of a joint downstream target node, with their joint effect on the target governed by an *OR* logical operator, this may constitute a synergy that is highly likely to stand out in an analysis, since the OR operator would cause either drug alone to not affect a joint downstream node. Between these two extremes, the topology will be more or less likely to produce a particular synergy prediction for a given combination perturbation. In order to correct for topology-intrinsic propensities for predicting some synergies we next employed a normalization strategy where synergy predictions for an automatically parameterized model ensemble are normalized to a *randomly* parameterized model, meaning a model ensemble that covers many different selections of OR and AND operators, irrespective of any particular stable state. This means that in our further analyses we used the fold change of the predicted synergy score of a test model against a randomly yet proliferative parameterized model (see Methods).

We first tested our software pipeline by considering predictions from model ensembles for simulating the 21 drug combinations that were analyzed previously^17^. Synergies were defined as a predicted ‘growth’ output for two drugs together being lower than the product of each individual drug’s ‘growth’, analogous to the Bliss synergy metric^30^ used in cell culture lab experiments (see Methods). Among 21 drug pairs, 15 were predicted to act synergistically by this definition, exhibiting a range of synergy strengths, and quantified performance of these models was surprisingly high: by selecting different thresholds for synergy predictions a receiver-operating characteristic (ROC) curve (sensitivity vs 1-specificity) shows a ROC area-under-curve (AUC) of 0.97 and a precision-recall (PR) AUC of 0.91, see figure 2C. The analysis shows that the top six predictions comprise all four experimentally validated synergies. Notably, the automatically parameterized models produced no false negatives. This effectively means we could in principle have reduced our full drug screen to only test 29% percent of the combinations (6 out of 21), had we guided our experiments by model simulations, which is comparable to the performance in our manually parameterized model^17^.

**Figure 2C:**
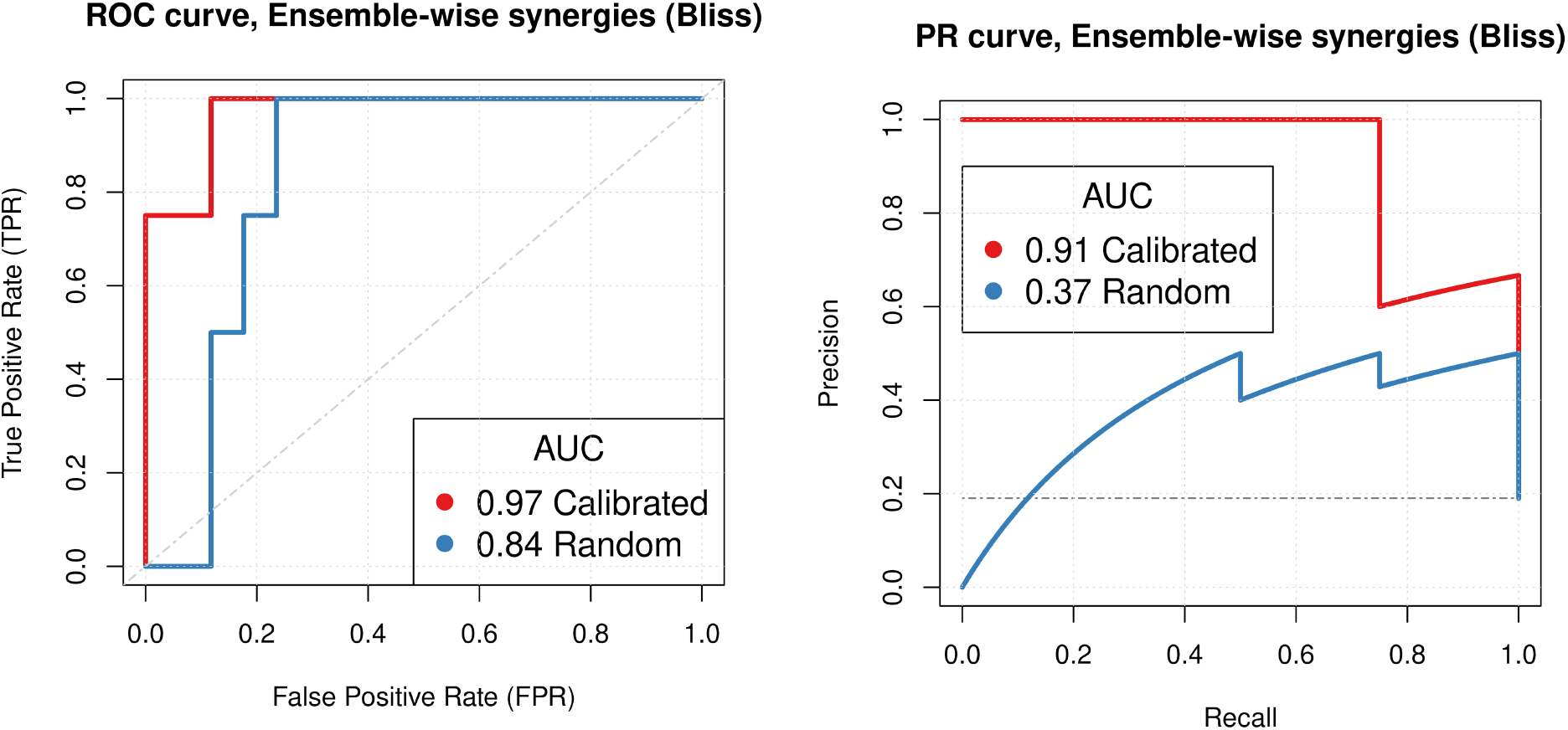
Predictive performance of ensembles of logical models. ROC curves are in the left panel, and PR curves are in the right panel. Random model predictions were generated by collecting predictions from ensembles of models trained to a random yet proliferative phenotype. Calibrated predictions were generated by model ensembles, trained to steady state data, and normalized to the random model predictions (see Methods for more details). The genetic algorithm modified the balance of influence between positive and negative regulators of a target node, while topology features (edges, nodes) were not modified. Both ROC and PR curves show very good performance across all model sets for the calibrated models, similar to results from Flobak et al (2015).

### Automated model optimization as a solution for larger model topologies

The benefits of automatic parameterization become more apparent in calibration of models with larger topologies. We demonstrate this with the CASCADE 2.0 model, which is a manually curated cancer signaling topology comprising 144 nodes and 367 edges. CASCADE 2.0 includes pathways with TGF-beta, JAK-STAT, and Rho GTPases, as well as extensions of pathways already present in CASCADE 1.0, to enable simulation of a larger set of drug combinations^18^. Starting with this large curated model, we analysed in more detail the effects of automated model training while randomly mutating logical rule configurations and network connectivity, and assessed the results against the biological regulatory mechanisms that were affected. We varied several aspects in the training protocol, each time assessing the effect on the performance of the models for correct synergy prediction:

- Optimising logical rules against partially incorrect calibration data
- Optimising the regulatory network by stepwise, random removal/inclusion of edges
- Checking the effect of random rewiring of the regulatory network

For each of these model alteration strategies, we not only looked for overall fitness but also in more detail at the represented biological mechanisms that were affected, to judge whether improved or reduced simulation performance could be reconciled with involvement of proteins of regulatory interactions in the context of cancer. The hypothesis was that, while taking the overall value of curated prior knowledge as a given, the relevance of individual regulatory interactions and the precise mathematical representation of their regulatory effects in specific cancer environments might be difficult to infer from papers and therefore could be algorithmically improved. The effects of the network connectivity and rule mutations were judged in model ensembles and compared with observed synergies. For each model able to reach a stable state, mutations also allowed to assess mutual dependencies between edges or subsets of edges and corresponding rules, possibly indicating context dependence. This allowed us to identify parameters and edges that appeared to be essentially fixed and thereby of fundamental importance for model performance.

We compared simulation results from automatically trained ensembles of 450 models to drug synergies in a new drug screen of the AGS cancer cell line comprising 153 combinations of 18 targeted drugs^31^. We found that, in contrast to the CASCADE 1.0 predictions, normalization of topology-intrinsic prediction propensities was critical to the predictive performance (see Supplementary Figures 1 and 2). We find that models obtained by automated optimization, as described above, could predict drug synergies with a ROC AUC of 0.69 and a PR AUC of 0.18, clearly outperforming random model predictions (see Figure 3). This means that our drug screen could have been reduced from blindly testing all 153 combinations to only the screening of 36 combinations, which would increase the synergy prevalence of tested combinations from 4% (6 of 153) to 11% (4 of 36). We would dismiss 117 drug combinations but at the expense of missing two observed synergies.

**Figure 3:**
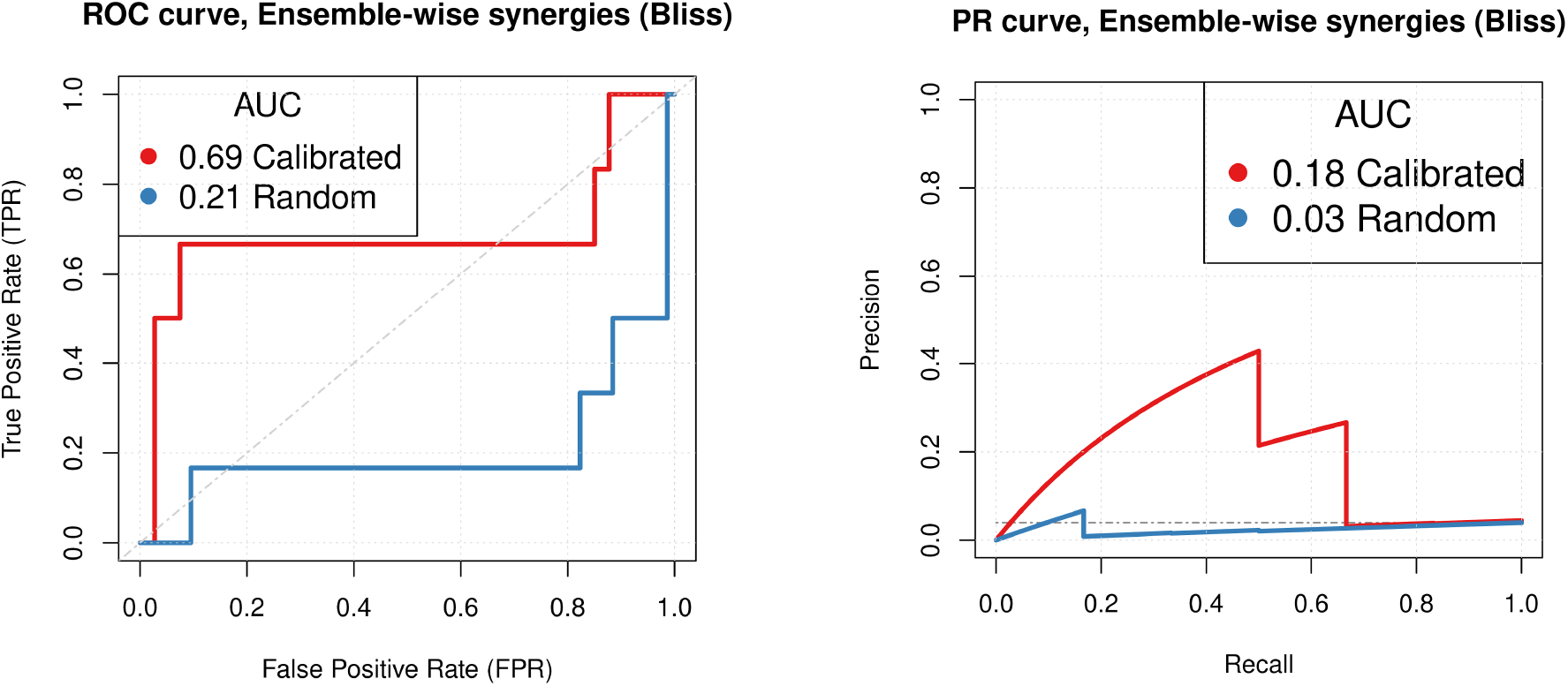
Model performance. Predictive performance of calibrated and random models based on the CASCADE 2.0 model was tested against data from a corresponding drug screen^31^. Models have logical rule mutations only and Bliss Independence was used to assess the model performance. We observe that correctly calibrated models perform substantially better than random models.

### Topology and calibration data needs to be correct

From here on we focus on the CASCADE 2.0 topology, for additional analyses see Supplementary Material, which has similar experiments with the CASCADE 1.0 topology, underpinning conclusions analogous to those drawn here.

#### Impact of data quality on model performance

Since our models are derived from prior knowledge and calibrated based on sample-specific measurements (calibration data), modifications to both the prior knowledge and data must be expected to affect the predictive performance of the models. We first checked how the quality of the calibration data affected model predictions. The performance of models trained to partially incorrect calibration data was expressed as PR AUC and, when plotted against the fitness of these models compared to the fully correct calibration data, we observe that higher PR AUC correlates with higher fitness of models, indicating that calibration of models indeed improved synergy predictions for our dataset (see figure 4). However, even models trained to highly incorrect data, with roughly 50% of calibration data flipped (meaning a true fitness around 0.5), perform better than a random classifier (PR AUC 0.04), indicating that model topology alone already carries information that can be leveraged to predict drug synergies. Note that due to the stochasticity of model calibration, the model ensemble average fitness never reached the extreme fitnesses below ~0.3.

**Figure 4:**
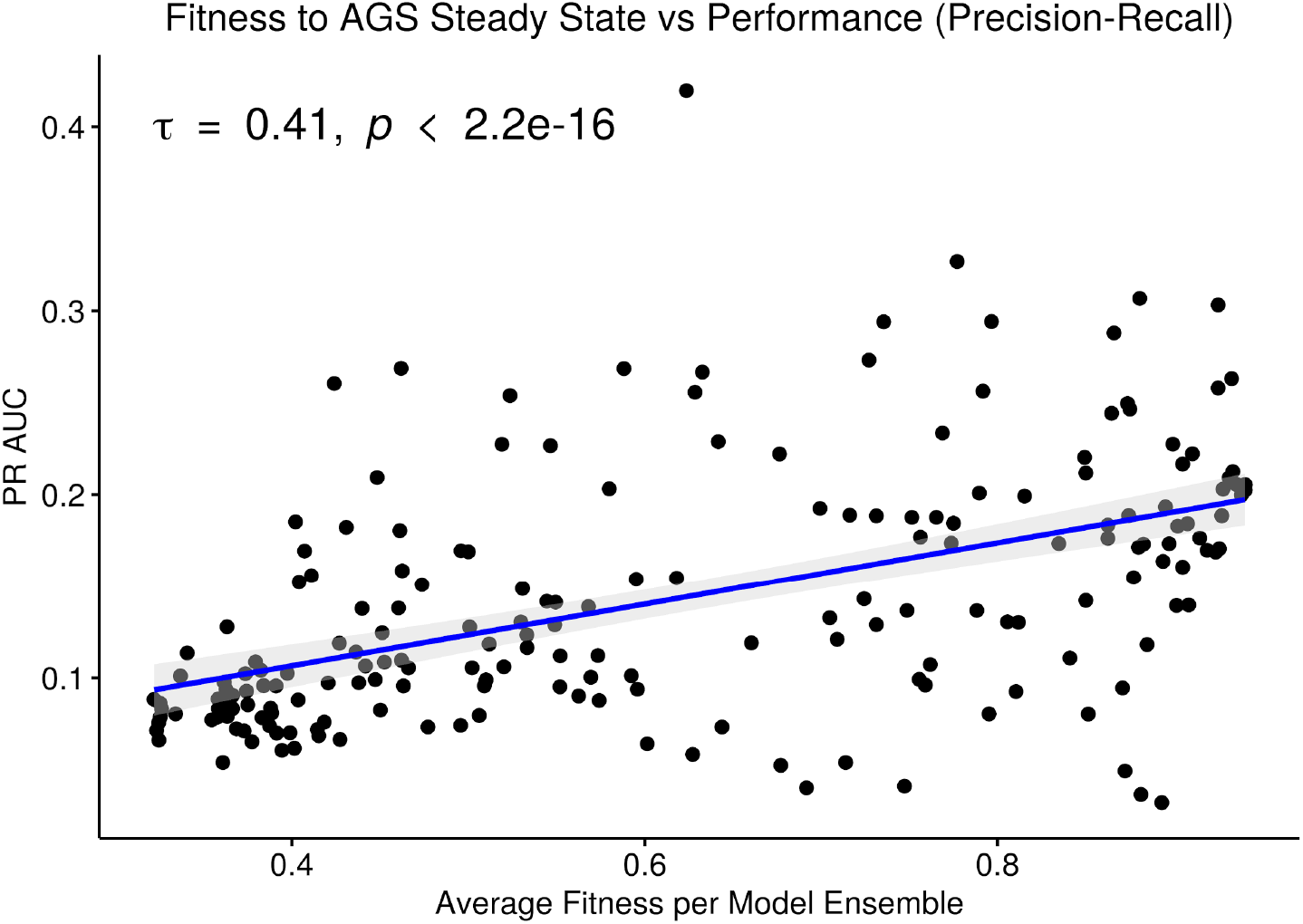
PR AUC performance dependence on fitness. Each model ensemble, displayed as one dot in the scatterplot, was trained to a partially incorrect steady state signaling profile derived from the biological phenotype of the AGS cell line^17^. A total of 205 training profiles were created, each one used to generate one model ensemble consisting of 60 models. The x-axis reports the average fitness of each model ensemble as evaluated to the curated steady state. Because of the non-normality of the data, the Kendall rank-based correlation^32^ test is used to derive the proposed association.

#### Randomizing regulatory edges of the curated model reduces predictiveness

As the quality of the calibration data does impact model performance, but not obliviate it even if these data are highly incorrect, we next explored the value of the quality of the curated regulatory graph topology. We generated a series of models with various degrees of ‘scrambled’ topologies with modified causal interactions in the regulatory graph (of the type: source - effect - target), by randomly exchanging a particular ‘source’ in a causal interaction with the source of another interaction and investigated the performance of models with these incorrect regulatory interactions. Similarly, we investigated the impact of randomly reassigning target nodes, as well as the impact of inverting signed effects, i.e. from inhibition to activation, and vice versa. Note that we will later explore the effect of missing prior knowledge (simple deletions), while here we present results for incorrect prior knowledge. The results (Figure 5) show that even low levels of randomization in the curated knowledge significantly reduce the predictive power of the models, quickly approaching random performance. Overall, we conclude that both calibration data and prior knowledge quality are important to correctly predict drug synergies, and that errors can be detrimental.

**Figure 5:**
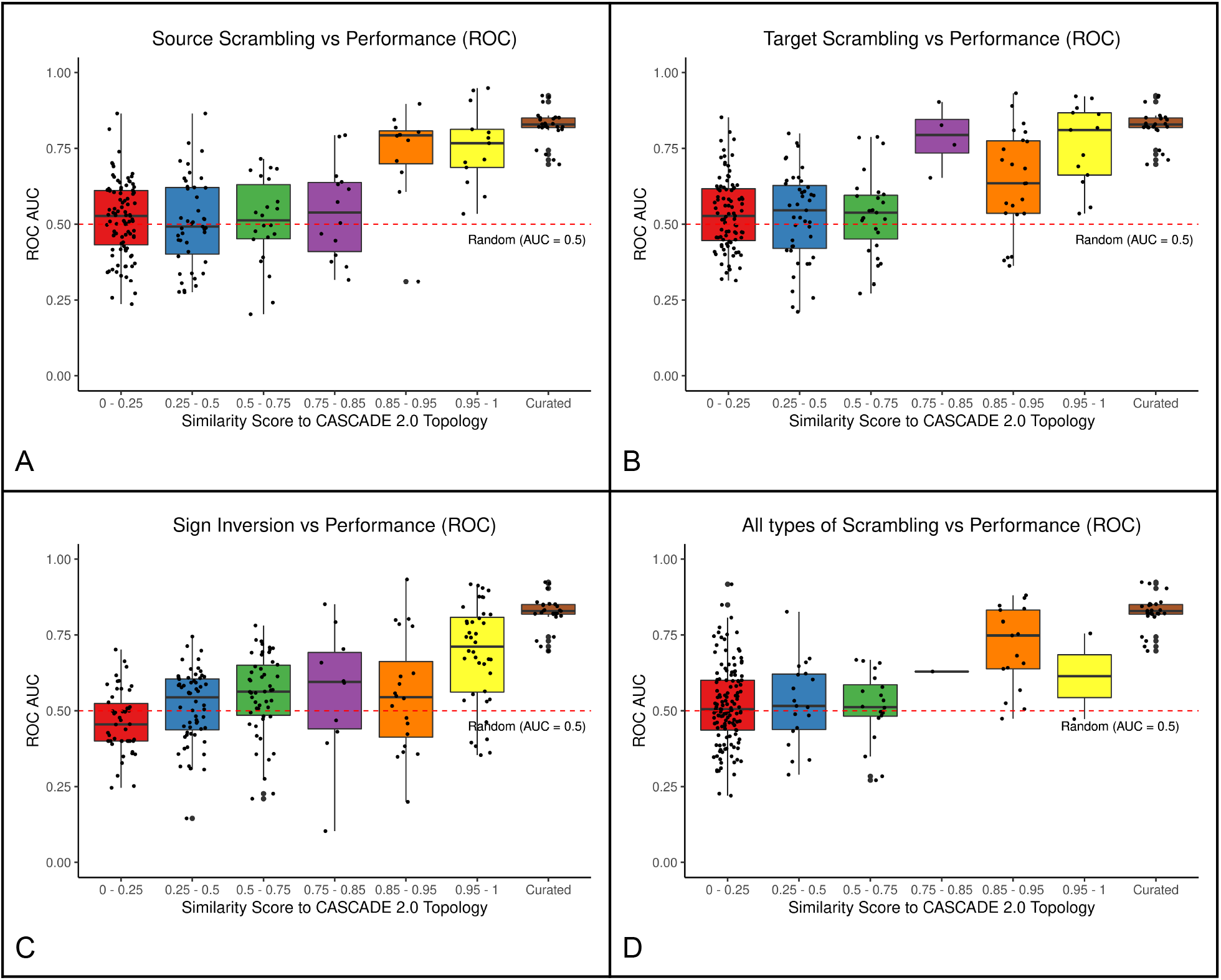
Effects of variations introduced in the CASCADE 2.0 prior knowledge graph. Panel A shows the effect of reassigning source nodes in causal interactions, panel B shows the effect of randomly reassigning target nodes in causal interactions, panel C shows the effect of randomly inverting activation/inhibition annotation, and panel D shows the result for all types of modifications introduced simultaneously. Each box plot shows a graded response for the predictive performance from complete modifications (left) to less substantial modifications (right). Each dot represents a different model ensemble generated from the associated topology, calibrated to the AGS steady state, and normalized to a random yet proliferative profile. The “Curated” group refers to model ensembles bootstrapped from a pool of models generated using our optimization algorithm from the original CASCADE 2.0 topology. See Supplementary Material Figure S4 for a similar analysis with precision-recall as performance metric, and Figures S5 and S6 for the same analysis done on the CASCADE 1.0 topology.

#### Model ensemble heterogeneity and mechanistic insight

To appreciate the heterogeneity amongst models in model ensembles obtained through the parameter optimization, we studied both attractor and parameterization heterogeneity against model fitness, in subsets of these ensembles selected for specific features (model sub-ensembles). In the heatmap grouped by K-means clustering (Figure 6), calibrated models to a relatively large extent obey the calibration data, with states of steady state nodes mostly identical to the data to which they were trained (the subset of nodes (24 of 144) that were specified in the calibration data). Model stable state vectors (rows in Figure 6) have notable areas of homogeneity, as judged by large stretches of nodes (indicated on the X-axis) that are either all activated (green) or inhibited (red) in all models, but in other areas (e.g. the upper-middle panel of the heatmap) the heterogeneity and discrepancies with calibration data is quite substantial. This heterogeneity was much more widespread in the parameterization space. For some nodes there is high correlation between their parameterization (link operator AND-NOT vs OR-NOT) and stable state (Inactive or Active, respectively), but for many the correlation is surprisingly low (see Agreement panel in Figure 6). These observations indicate that a) a limited set of training nodes was sufficient to provide homogeneity in parts of the attractor space, and b) large heterogeneity in the parameterization space still can be compliant with homogeneity in the attractor space (see in particular the large green (active) area of the stable-state heatmap). In other words: there are many logical rule configurations that yield models properly representing biological observations compliant with calibration data. This underpins the decision to use model ensembles rather than single models, since these ensembles cover a larger set of parameterizations (behavior) that are all compliant with the input data.

**Figure 6:**
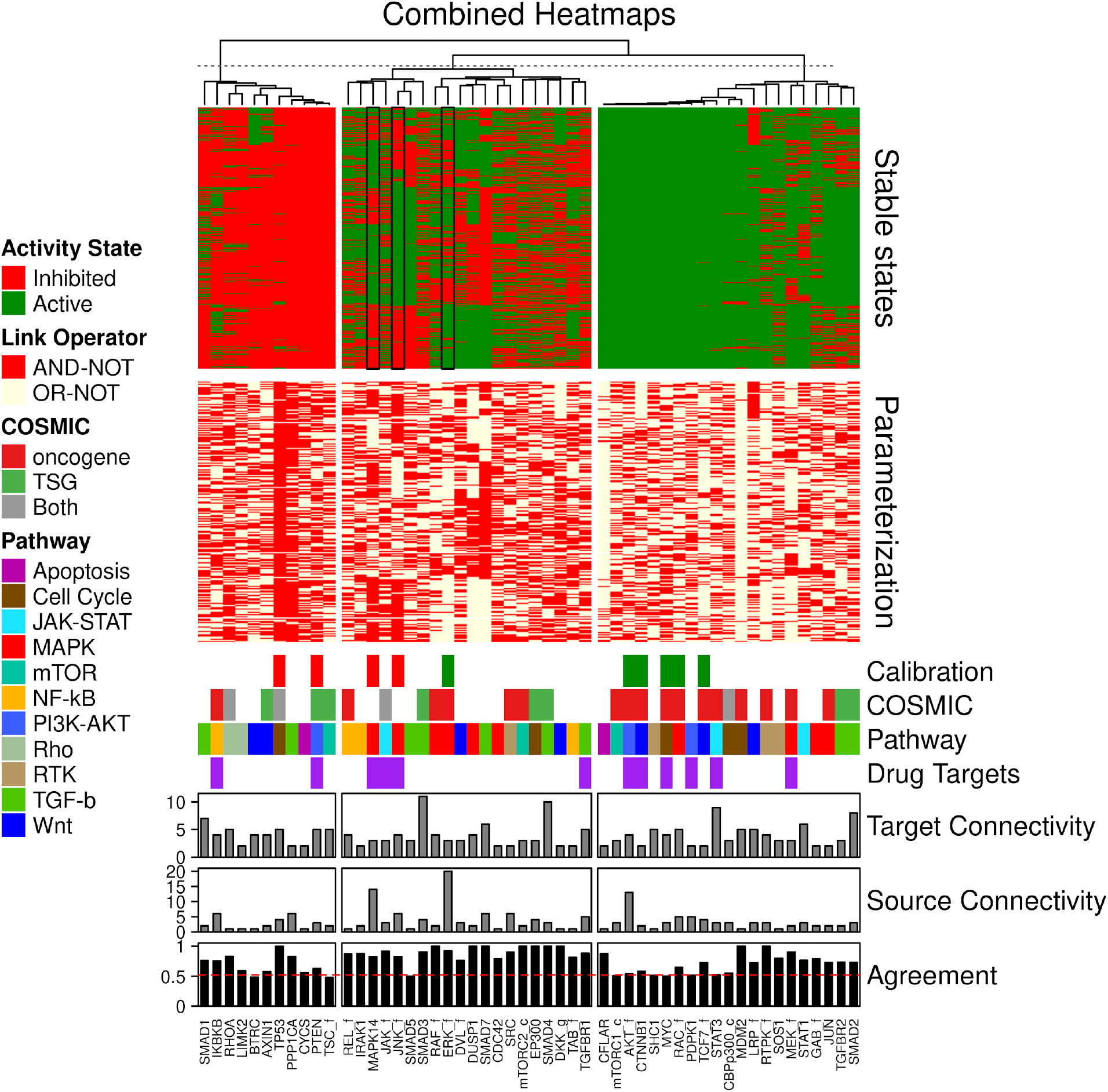
Combined stable states and parameterization heatmaps. A total of 4500 Boolean models were used for this analysis. Only the CASCADE 2.0 nodes that have a link-operator in their respective Boolean equation are shown. The 52 link-operator nodes have been assigned to 3 clusters with K-means using the stable states matrix data. The link-operator data heatmap has the same row order as the stable states heatmap. Steady state data (Calibration), COSMIC classification of tumor suppressor genes (TSG) and oncogenes, pathway association (Pathway), drug target characteristic (Drug Targets), in-degree connectivity (Target connectivity), out-degree connectivity (Source connectivity) and percent agreement between parameterization and stable state annotations (Agreement) are indicated below the heatmaps.

The analysis of node states and parameterization allows us to investigate mechanisms underlying observed behavior and to look for biological explanations for some of the observations. As shown in Figure 6, indicated by panel ‘COSMIC’, the models allow the prediction of activity of several proteins implicated in cancer. Figure 7 shows the analysis of their activity state, and it appears that proteins from genes annotated as oncogenes in COSMIC tend to be active, while proteins from genes annotated as tumor suppressor genes (TSG) tend to be inactive in overall steady states. This is biologically plausible and attests to the capability of our mechanistic model to generate hypotheses about the underlying molecular biology.

**Figure 7:**
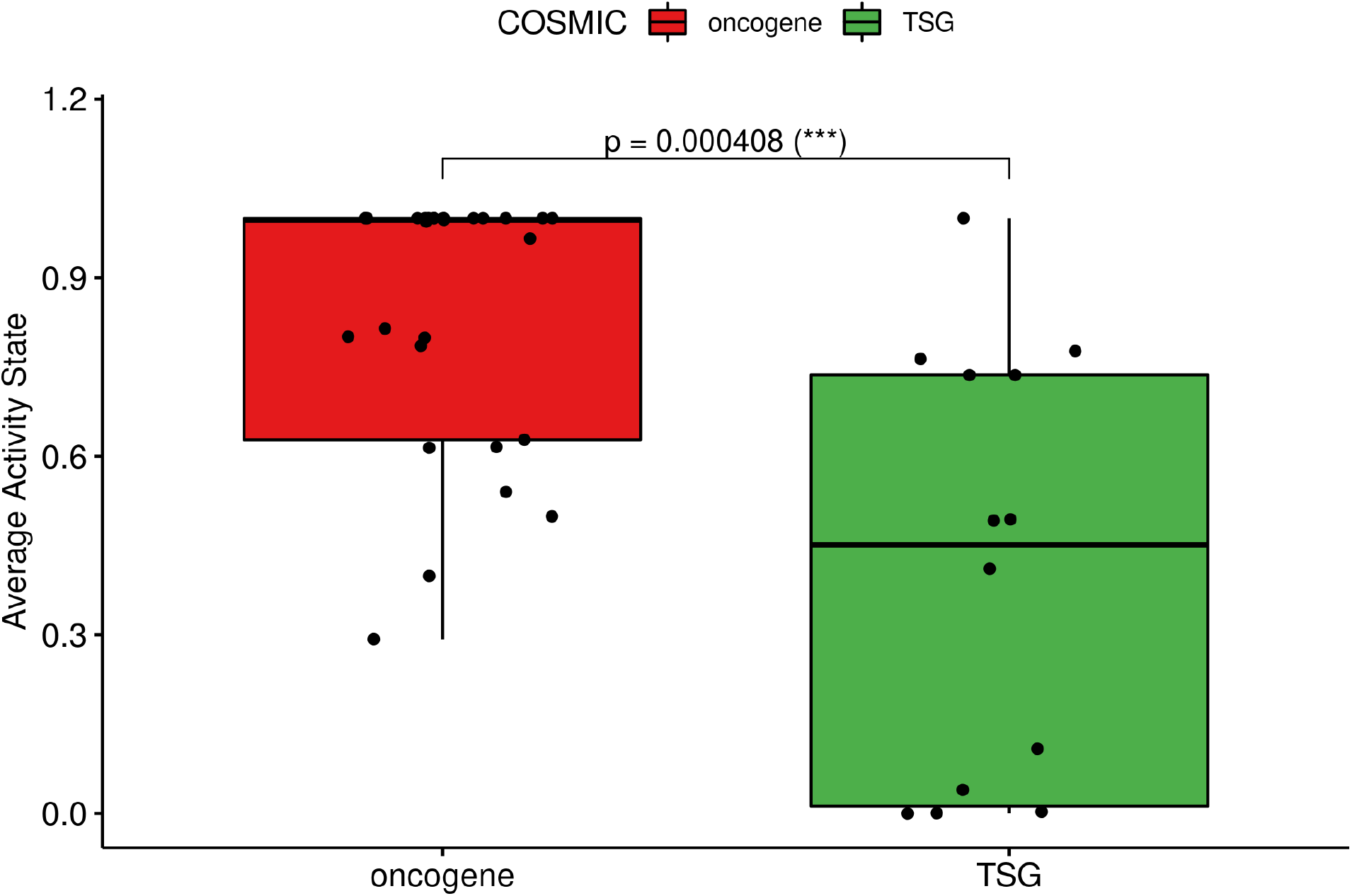
Box-plot of stable state protein activities. The stable state models yield activity values for all proteins and these activities are displayed for oncogene and tumour suppressor gene proteins. Proteins from oncogenes (left) tend to be designated as active, while proteins from tumor suppressor genes (TSG, right) tend to be designated inactive.

The training of the models to biomarker data never results in the absolute maximum fitness (1), and the stable state analysis (Figure 6) shows that three data points are most often violated: JNK signaling (JNK_f), ERK signaling (ERK_f) and p38 signaling (MAPK14), all member of the MAPK pathway (see Figure 6, stable state panel, black rectangles). These network nodes are all clustered in the highly heterogeneous section in the steady state heatmap. In the manual curation of the CASCADE 1.0 topology^17^ it was noted that reports on the activity of ERK in AGS cells varied frequently, with only slightly more than half of the publications reporting ERK to be active. We found that the predictive performance of model versions with ERK being active was significantly higher than the sub-ensemble where ERK is inactive (see Figure 8), suggesting that from a functional point of view ERK should be considered active in AGS cells.

**Figure 8:**
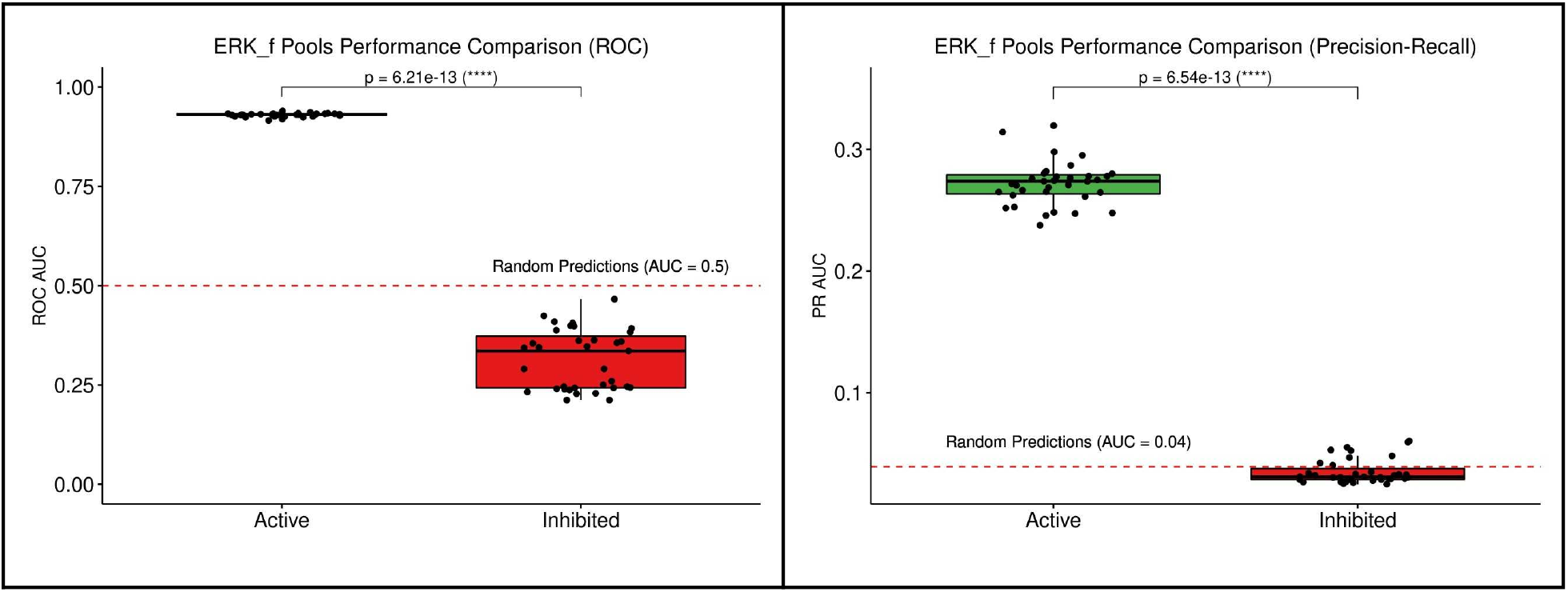
Comparison of predictive performance of model sub-ensembles. The model set with ERK active scores better than the models with ERK inactive, both for ROC (left panel) and PR (right panel).

### Rule and edge optimization identifies key regulatory mechanisms

Traditionally, logical model definitions start out with a prior knowledge graph, after which a most optimal parameterization is sought based on experimental evidence and model behavior. We asked which was most influential to accurately predict synergies: alterations to the topology or to the parameterization, in the evolution to maximum fitness. We configured the genetic algorithm to modify edges in the topology, by either removing or subsequently restoring edges from the initial prior knowledge network (as long as no signaling component lost all its source inputs), or to modify the parameterization of logical rules. We found that modifications of the edges and the parameterization both resulted in substantially improved prediction performance, significantly better than a random classifier. In particular, the performance (as evaluated by precision-recall) was very high for topology-mutated models, even outperforming models trained by parameterization optimization. While ROC AUC was consistently high around 0.8, the PR curve benefited by displaying very high positive predictive values at very conservative sensitivity thresholds, meaning that a predicted drug synergy is highly likely to represent an actual synergy, see Figure 9. Note that there is a major difference between missing data and incorrect data: as was previously demonstrated (Figure 5), model predictive performance suffered severely from included and incorrect prior knowledge, while model predictive performance can improve by omitting putatively correct prior knowledge.

**Figure 9:**
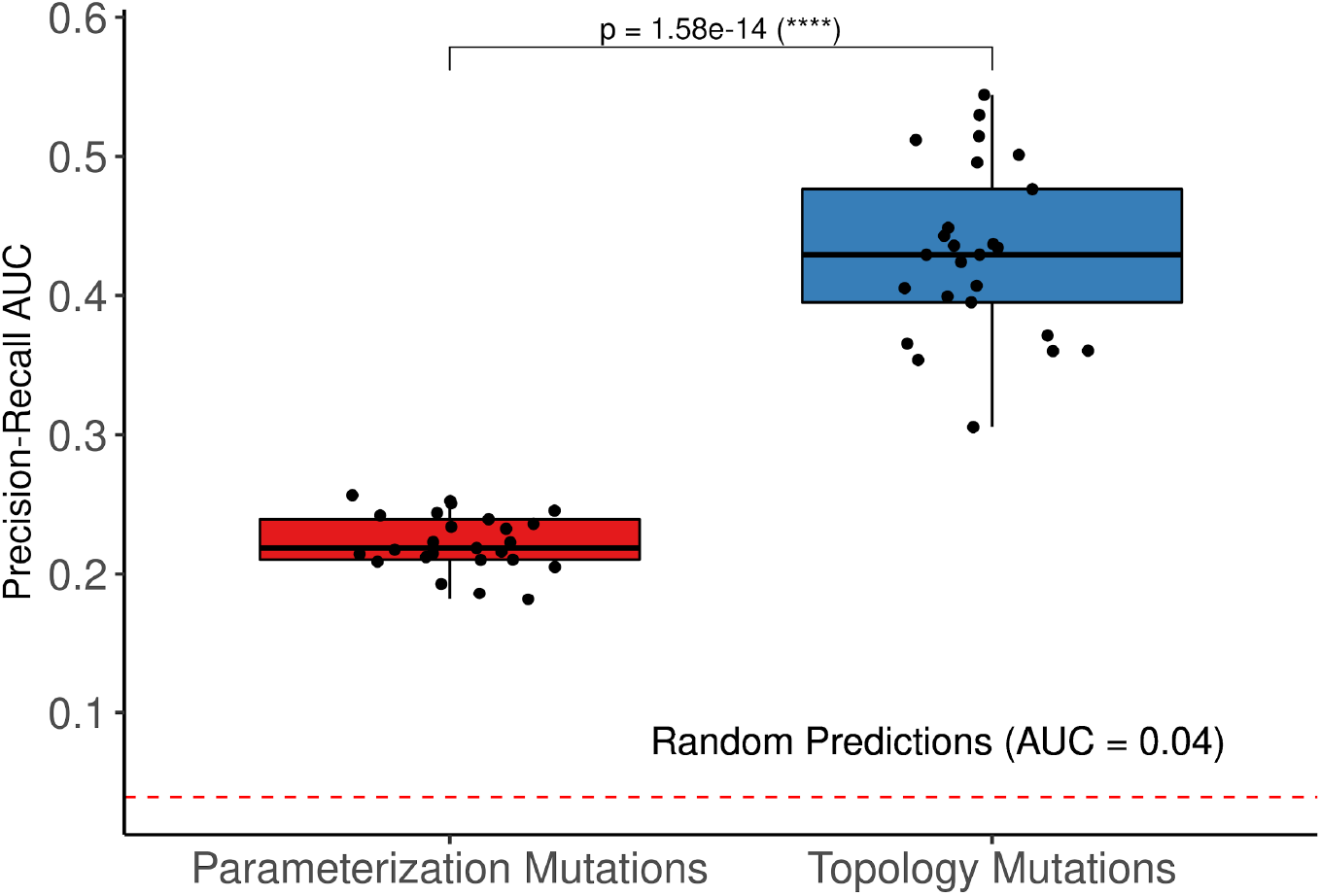
Model performance after parameter and topology modifications. Mutating parameterization (left) and topology (right) both tend to improve synergy prediction performance, as evaluated by precision-recall AUC. Models with topology modifications perform better than models with parameterization modifications.

Looking at the topology modifications, we hypothesized that deletion of certain edges could be favored by the genetic algorithm, to obtain maximum fitness. Every node in CASCADE 2.0 is annotated to a specific pathway^18^, allowing us to assign all edges to a specific pathway, if both source and target node belong to the same pathway, or to crosstalk for edges that link nodes from different pathways, see figure 10. We observed that certain edge groups are always preserved (the left-most cluster), while other edges are very likely to be removed (cluster on the right). Interestingly, a majority of these removed edges belong to the TGF-beta pathway, in particular representing inhibitory effects of the protein SKI and other inhibitors of SMADs. In the model, SKI itself is inhibited by active AKT signaling, and thus removal of inhibitory edges from SKI allows restricting the activity of some of the SMAD proteins in the model, in particular to SMAD1, SMAD3 and SMAD4, which tend to be inactivated in the topology mutated models. It is evident from figure 10 that crosstalk is largely preserved during model optimization, potentially relating not only to sparse knowledge of crosstalk in the prior knowledge network, but also to the biological importance of signaling that is not confined to what was more or less arbitrarily viewed as pathways.

**Figure 10:**
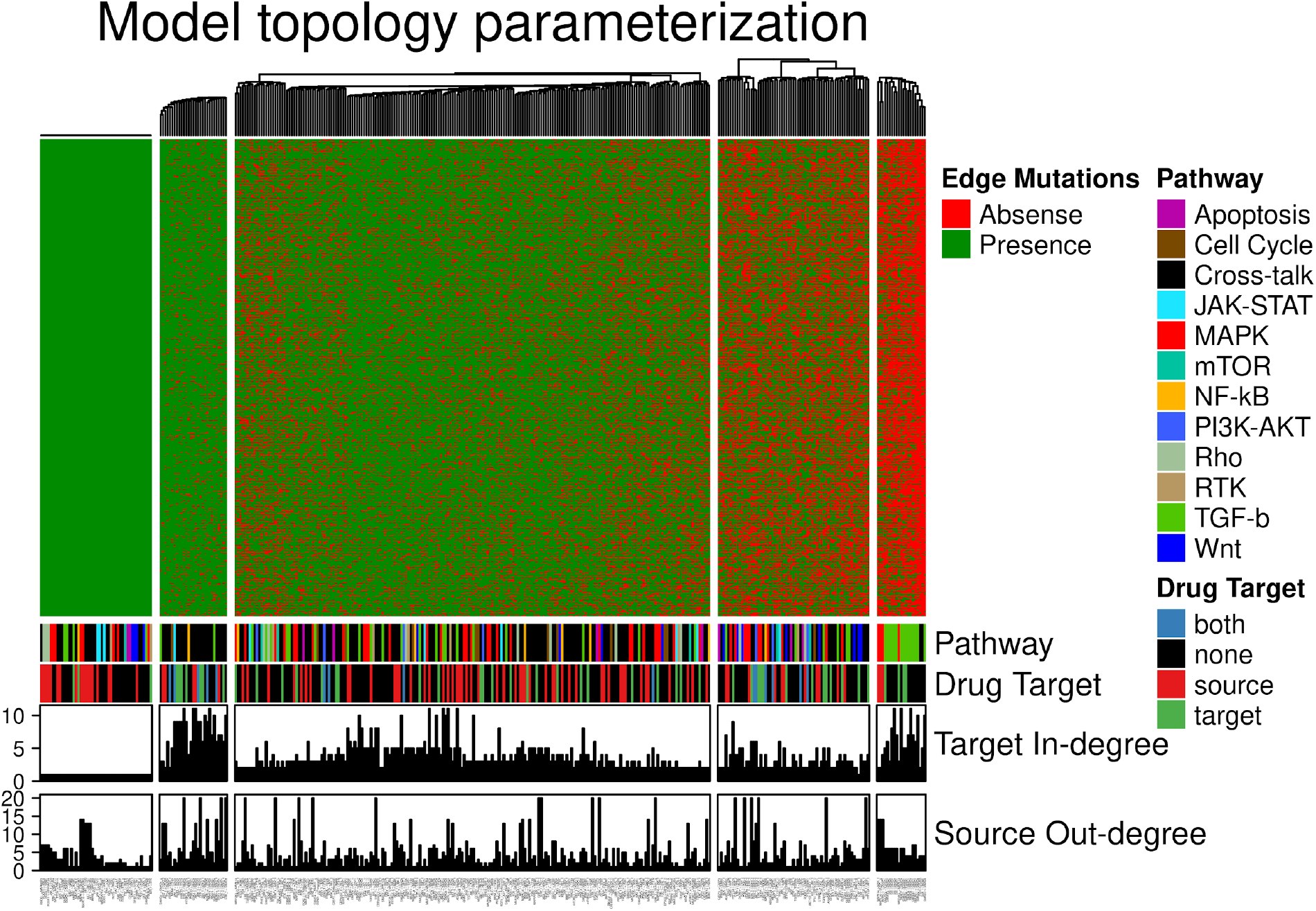
Heatmap of steady state models with topology mutations. Each column represents an interaction, each row represents one model, all rows jointly represent the ensemble. Five clusters, guided by K-means clustering, stand out: the first cluster from the left represents edges that cannot be removed because nodes would lose regulations. The second cluster from left represents nodes that are likely to be preserved, the middle cluster represents edges that are often preserved, the fourth cluster from left represents edges that are often discarded, and the last cluster, right, represents edges that are almost always discarded in the evolution to maximum fitness.

### Validation of synergy predictions in vivo

In order to test the translational relevance of our drug synergy prediction platform we performed in vivo validation for one of the proposed novel drug synergies. The synergies of TAK1 inhibition combined with PI3K inhibition, already identified in our previous logical modeling work, had not been reported earlier and thus represent novel synergies of potential interest in future cancer therapy. We rediscover the same synergy of combined TAK1 and PI3K inhibition in our framework for automated model parameterization.

In order to test reproducibility across different high throughput drug screen readouts we subjected AGS cancer cells to combined TAK1 and PI3K inhibition and monitored the response both by ATP content measurement (viability) and by microscopy (confluence), see Figure 11. For both readouts there is a region of synergistic response to drugs applied at medium doses as indicated by the drug concentration gradients. We subcutaneously injected AGS gastric adenocarcinoma xenograft tumors in Balb/c mice to test the synergy of combined inhibition of TAK1 (5Z-7-oxozeaenol) and PI3K (PI103) *in vivo.* The xenograft tumors (n=30) were randomized to four groups: control, PI103, (5Z)-7-oxozeaenol and a combination group which received both PI103 and (5Z)-7-oxozeaenol, see figure 12A. At the end of the experiment the combination group displayed significant changes (t-test) in relative tumor size compared to either single-treatment group (Figure 12B and 12C). We observed a similar reduced proliferative capacity for tumours in mice treated jointly with TAK1 and PI3K inhibitors, as indicated by tumour proliferation marker Ki67 (Figure 12D and 12E). The clear difference between the significant tumor growth inhibitory effect of the combination and the non-significant activity of individual agents strongly indicates a synergistic anti-tumor effect of the two agents. A concern for drug synergies is that possible side-effects might also be expressed in synergistic ways. We therefore chose doses to be at the lower end of effective concentrations, compared to previously published in vivo use of the inhibitors^33, 34, 35, 36^. Despite low dosage the inhibitors together reduced tumor growth, without signs of pain or weight loss.

**Figure 11:**
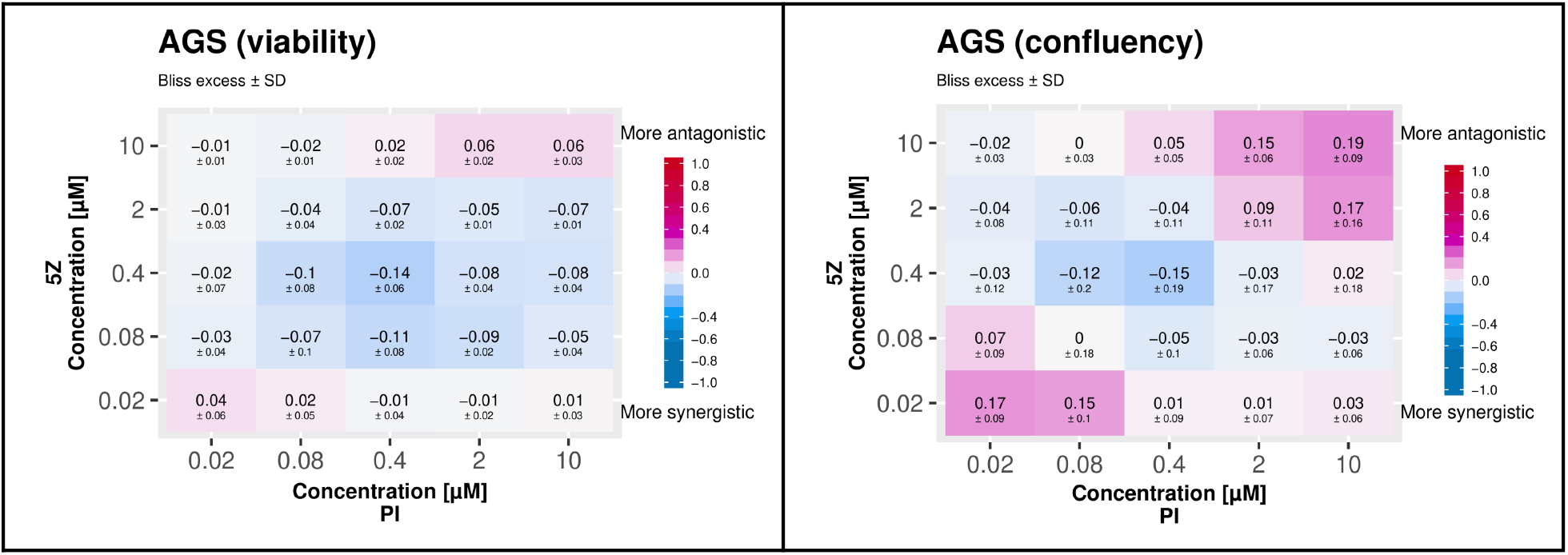
Synergy of the drugs targeting TAK1 (5Z) and PI3K (PI) confirmed in viability (left) and confluency (right) screens.

**Figure 12A:**
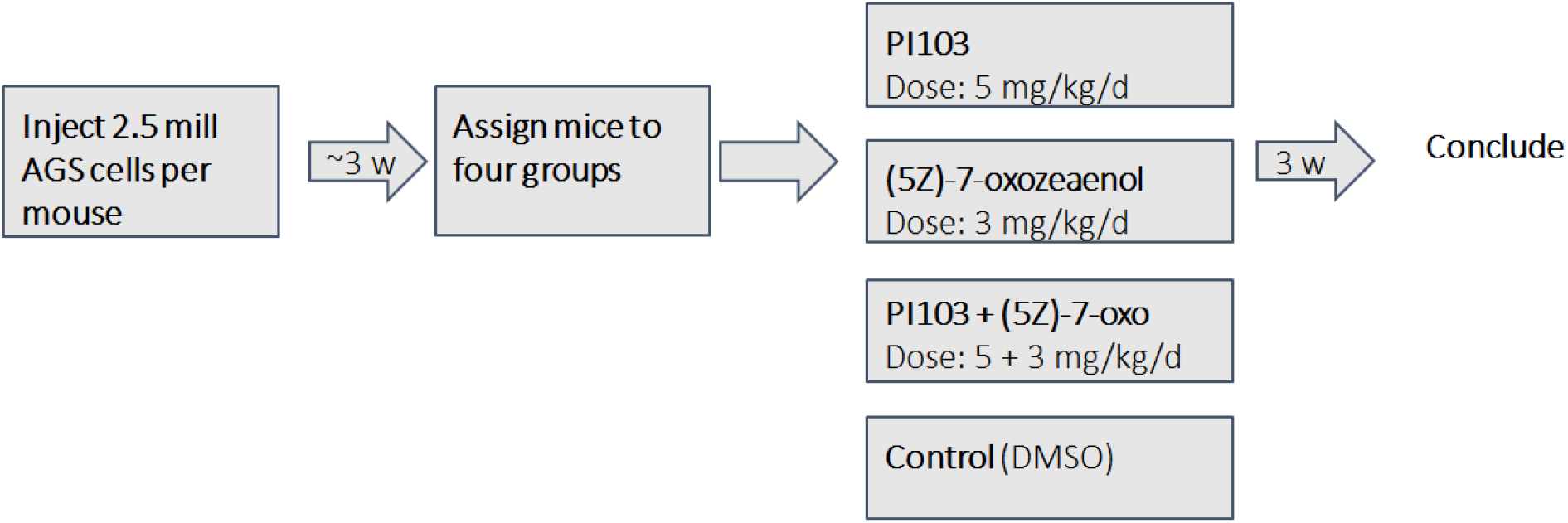
Xenograft experimental design. Cancer cells were injected subcutaneously in mice and allowed to grow until a visible tumour could be identified, after which the mice were randomized into four groups. Mice were then treated with single drugs and the combination for 19 days after which the experiment was stopped and tumor sizes evaluated.

**Figure 12B:**
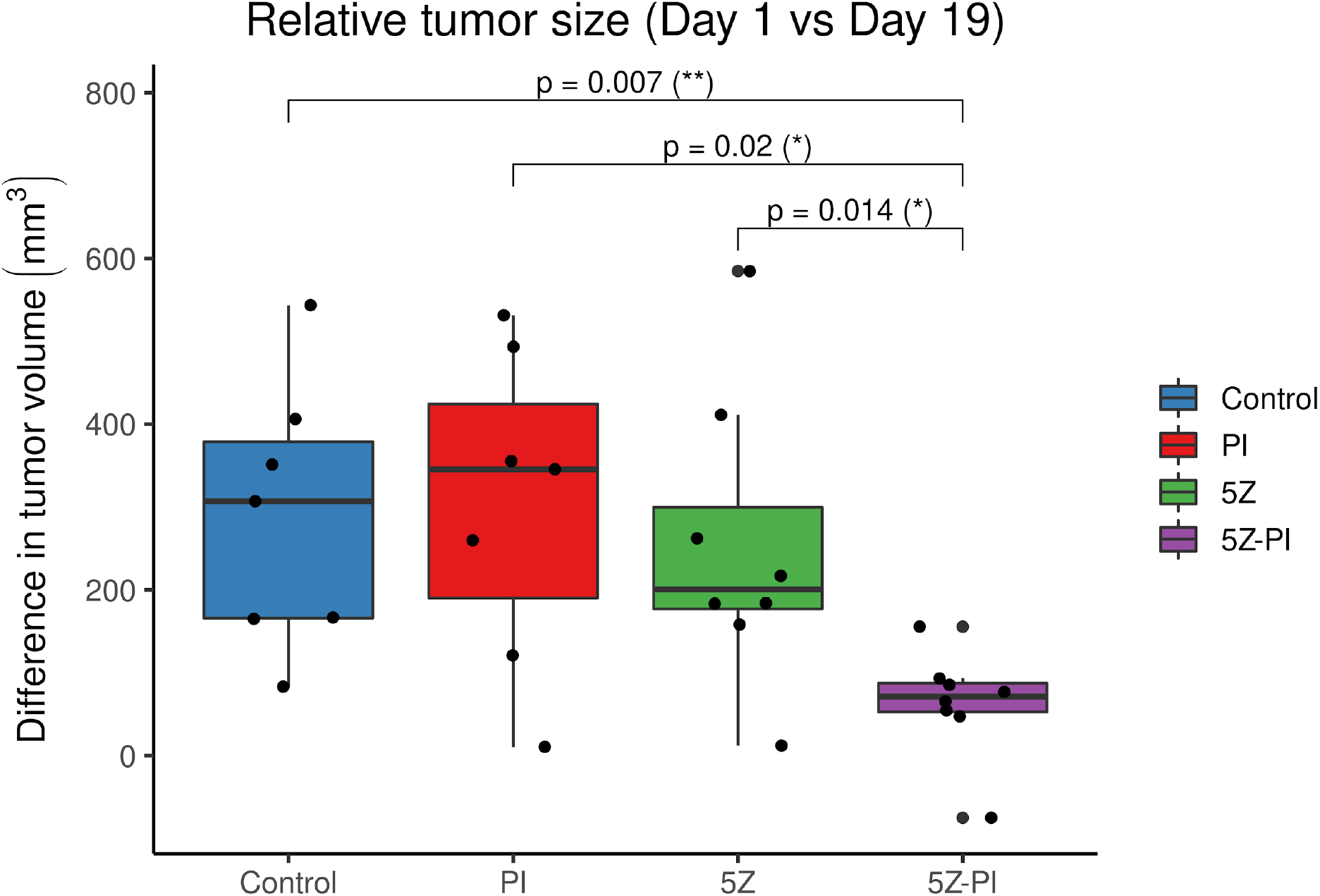
Testing of drug synergy in a mouse xenograft model. Tumor volumes determined at the end of the study were compared with tumor volumes at treatment onset. Tumors in mice receiving both inhibitors 5Z-7-oxozeaenol 3 mg/kg/d (5Z) and PI103 5 mg/kg/d (PI) show a smaller increase in size compared to either of the groups receiving only single inhibitors, and the control group. The combination effect was statistically significantly different from either single drug therapy as evaluated by Mann-Whitney U tests with corresponding p-values shown above the boxplots. Similar results were obtained using t-tests with a chosen significance level of p=0.05.

**Figure 12C:**
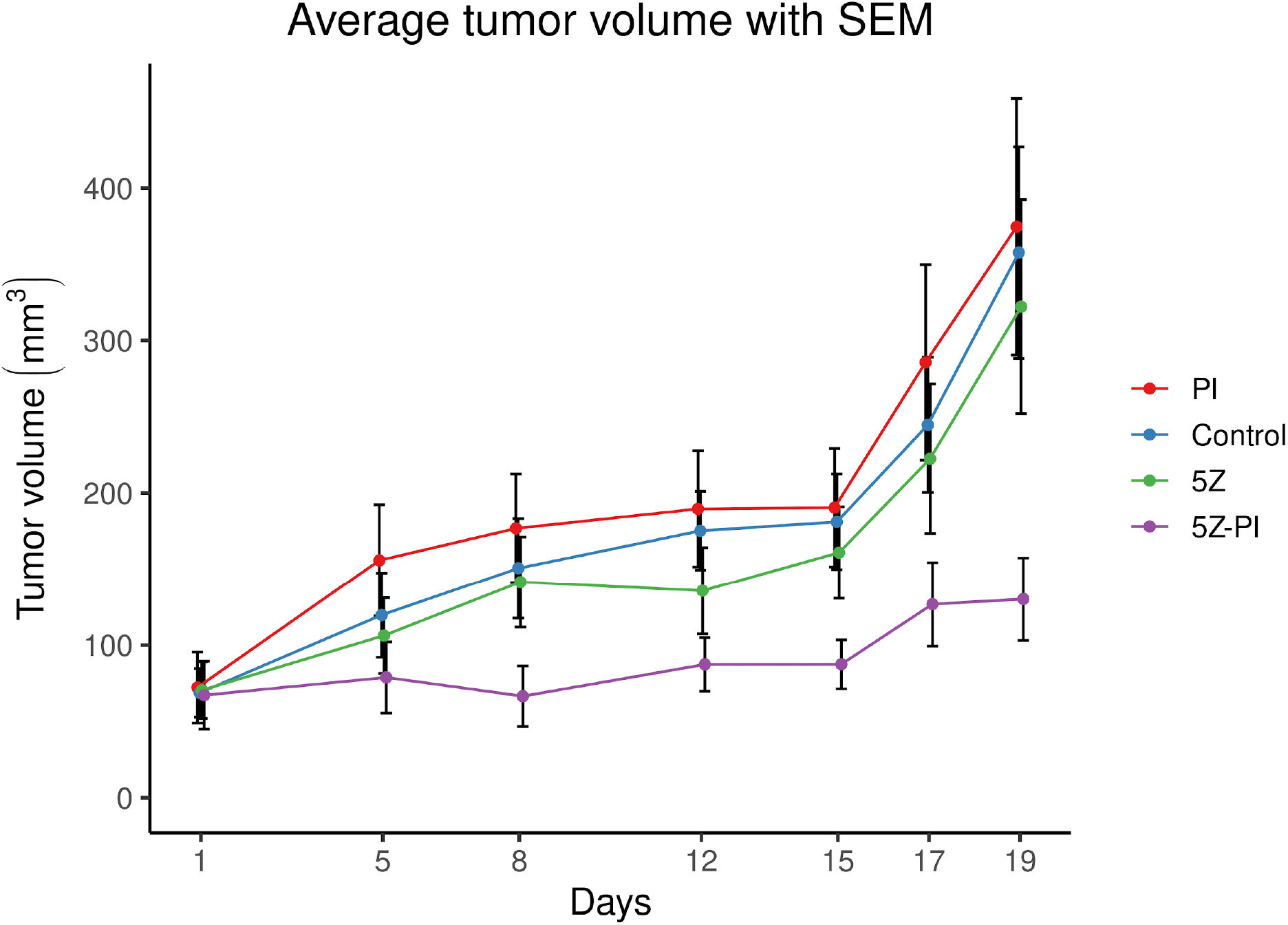
Average tumor volume for the four groups of mice with standard error of the mean (SEM) indicated by the error bars. The group receiving both inhibitors (5Z + PI) displays a more inhibited tumor growth than either of the groups receiving each single inhibitor and the control group.

**Figure 12D:**
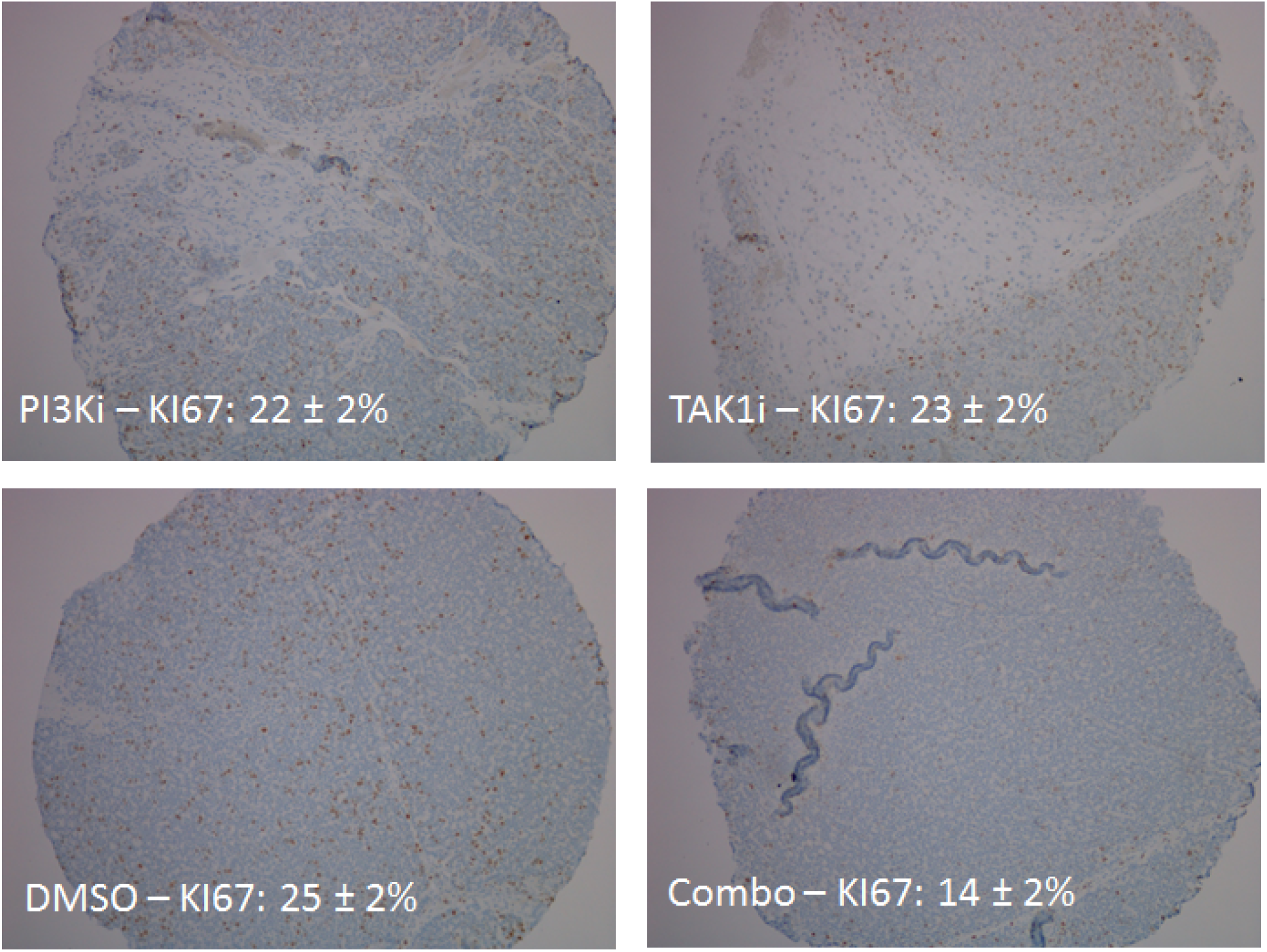
Ki67 proliferation index (count of Ki67 positive cells) reduced upon joint combination of TAK1 and PI3K inhibition.

## Discussion

Our results show that a curated prior knowledge network with an initial set of logical rules can be automatically parameterized by a genetic algorithm, using a fitness score reflecting how well the global stable state of a model matches the experimentally determined local states of signaling components of a cell line in its native growing state. We showed that the predictive performance from an automatically parameterized ensemble of models was on par with our original, curated CASCADE 1.0 model. We next used our parameterization software to calibrate the larger CASCADE 2.0 network topology. With larger topologies, benefits of automation become more apparent, and can be used to enable simulation for larger numbers of drugs and numbers of cell lines.

Finding drug synergies among the vast set of possible combinations of drugs calls for new approaches. To rationally reduce the prohibitively large experimental search space, we found that our approach can be highly useful to identify sets of drugs that are unlikely to display synergy and that should not be prioritized for testing. Even tackling the combinatorial complexity for standardized cancer models is already challenging, as exemplified by the AZS-DREAM Challenge^37^, where pairwise combinations of 118 drugs (6903 possible drug-drug combinations) are tested against a panel of 85 cell lines. If all combinations are screened in 6×6 matrices this corresponds to over 200.000 384-well plates for four technical replicates, clearly indicating that a trial-and-error approach is not economic for drug synergy discovery. Whereas most approaches for drug synergy predictions rely on perturbation data for training a classifier^38, 39, 40, 41^, our approach works well with calibration data based only on data from an unperturbed system, which greatly reduces the cost of data acquisition and opens possibilities for clinical applications.

Assessing the performance of drug synergy predictions is met with several challenges. First, targeted drugs, as those employed here (‘small molecule’ inhibitors), can affect a number of other targets in addition to their intended target, known as ‘off-target’ effects^42^. Since our simulation is based on canonical drug targets annotated for each drug, any information missing about off-target effects must be expected to impact simulations. In addition, drug synergy is an elusive concept itself, with different mathematical reference models producing different synergy scores^43^. Finally, high throughput drug screens, as employed here, typically reports drug responses based on measured residual ATP content after drug exposure, which is known not to capture all growth-reducing drug responses^44, 45, 46^. Despite these limitations, which must be expected to reduce the performance of any drug synergy prediction approach, our logical simulation-based in-silico pre-selection approach performs immensely better than a blinded screen that would assay the same numbers of candidates: at a sensitivity of 50%, roughly 35-40% of a pre-selected set of predicted synergies will be observed in follow up drug synergy experiments in drug screen where only 4% of drug combinations acted synergistically overall.

Our choice of a logical framework for computational simulations comes both with some benefits and limitations. Logic equations are very quick to evaluate, with high simulation speed enabling extensive simulations even on regular desktop computers. However, logic equations as employed here only allow two activity states for model components: active and inactive. Moreover, only two interaction strengths between components are allowed: full interaction or no interaction. These limitations, however, still to a large extent meet the demands and possibilities offered by experiments with present day laboratory techniques. Data from these experiments often lack finer-grained observations that would be needed for continuous modeling approaches and therefore logical modeling represents a valid compromise between molecular data richness and computation speed. Our implementation of logical model simulations only computes stable states, thereby discarding any potential complex attractor of models. This choice was based on computational efficiency, as computation of complex attractor in addition to stable states would severely tax our simulations. One possible avenue for future research will be to account for also complex attractors, either by approximations as offered by e.g. trap space analysis, or by a full characterization of model behaviour.

We find that our approach is somewhat sensitive to errors in the calibration data, and even more sensitive to errors in the prior knowledge, indicating that curation quality is paramount to our modeling approach. This demands for adequate causal statement curation protocols and standards that feed into high quality general and cancer signaling databases, a demand that is materializing^47, 48, 49, 50, 51^.

Interestingly, we found that deleting a small fraction of the regulatory links from the prior knowledge network can be very powerful in optimizing models for drug synergy predictions, whereas randomly rearranging regulatory links is very detrimental to model performance. Logical model construction can be performed by curating data resources or the literature, or by relying on high quality curated databases like Signor, SignaLink or IntAct. If quality of these resources, or the ad hoc curation of literature, would not be of high standard, this would significantly limit the performance of the resulting model. On the other hand, calling a prior knowledge network complete is essentially a judgement call, as all models are limited. This demonstrates that, while lacking in completeness, high quality curation can produce models that can predict drug synergies. Furthermore, whereas the inclusion of a regulatory link ideally needs evidence from observations about the functional relevance of such a link in the cell that is modeled, our observations about ERK activity highlight the variability of experimental data concerning such observations. It is therefore not unreasonable to accept that an optimization algorithm can choose to dismiss regulatory links for the benefit of improved model performance. The increasing availability of high quality curated molecular causal interaction data opens a perspective to fully automated model building, where algorithmic topology optimization can fine tune a model to perform adequately, for any target node requirements and cell type for which baseline biomarker data is available. A welcome feature of automatically parameterized logical models is their innate ability to suggest mechanisms underpinning a particular observation. Such model-driven hypotheses can lay foundations for targeted follow-up experiments that provide observations for directed model revisions, resulting in a model with higher validity.

It has been suggested that network topology alone already explains drug synergy^52, 53^, and that parameterization to a lesser extent defines or refines synergy predictions^54^. Experimentally it has been observed that drug synergies tend to vary between cell lines, with the most frequently observed synergistic drug pair only effective in about half of the cell lines analyzed^55,56^. In our analysis we observed that drug synergy predictions depend both on the specific parameterization of a given topology and on the topology itself. When we put our manually defined topology to the test, drug synergy predictions are more accurate for models optimized to represent cell-specific baseline biomarkers in their local states, compared to unconstrained local states. One may speculate that the interactions relevant to describe drug combination effects represent a subset of all potential (general) interactions, and that this subset varies from cell line to cell line. From a completeness perspective, given the limited knowledge of molecular biology today, any model representation will be a major simplification of reality, yet some of these models work.

## Author contributions

ÅF: Devised the project and developed the main conceptual ideas. Designed and developed prototype for simulation pipeline. Developed and implemented drug response combination simulation software. Analysed drug response prediction data and model performance. Designed and executed cell line experiments. Manually curated drug combination response data. Designed and executed xenograft experiment. Supervised drug combination matrix experiments. Supervised tissue microarray experiments.

JZ: Developed and implemented production-ready drug response combination simulation software. Executed various simulations, analysed drug response prediction data and model performance, produced resulting figures, produced supplementary information, produced github repositories, made all analyses reproducible.

MV: Analysed drug combination data. Developed and implemented drug response combination simulation software.

TS: Supervised xenograft experiments.

LT: Supervised xenograft experiments.

AG: Prepared and analysed tissue microarrays of xenograft tumours.

BN: Manually curated drug combination response data. Designed and executed drug combination matrix experiments.

EF: Manually curated drug combination response data. Designed and executed drug combination matrix experiments.

MK: Analysed drug response prediction data and model performance. Supervised the project. All authors contributed to the manuscript.

## Supplementary Figures

**Figure S1:**
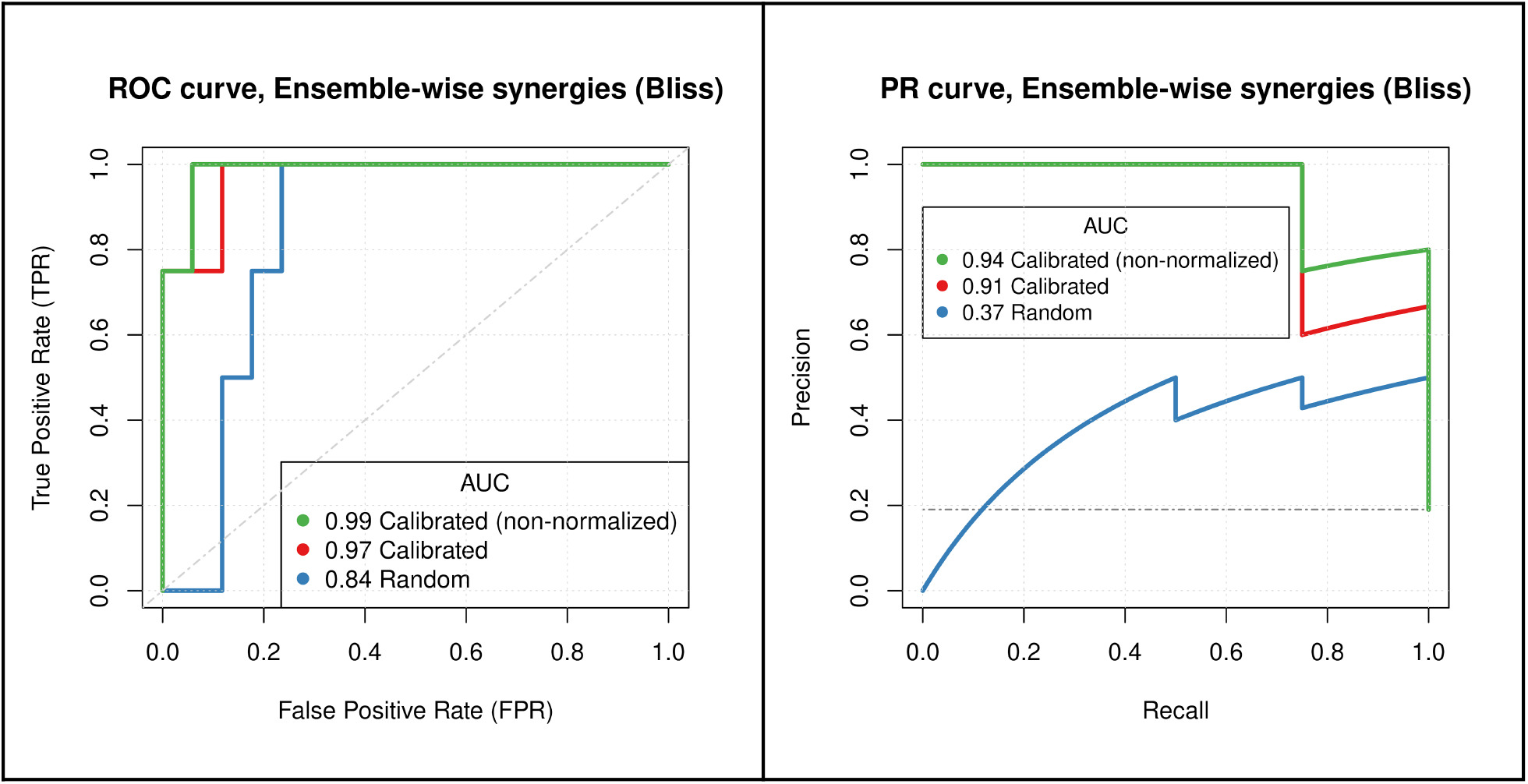
Predictive performance for random (proliferative) models, calibrated models (non-normalized) and calibrated normalized to random models (CASCADE 1.0 topology).

**Figure S2:**
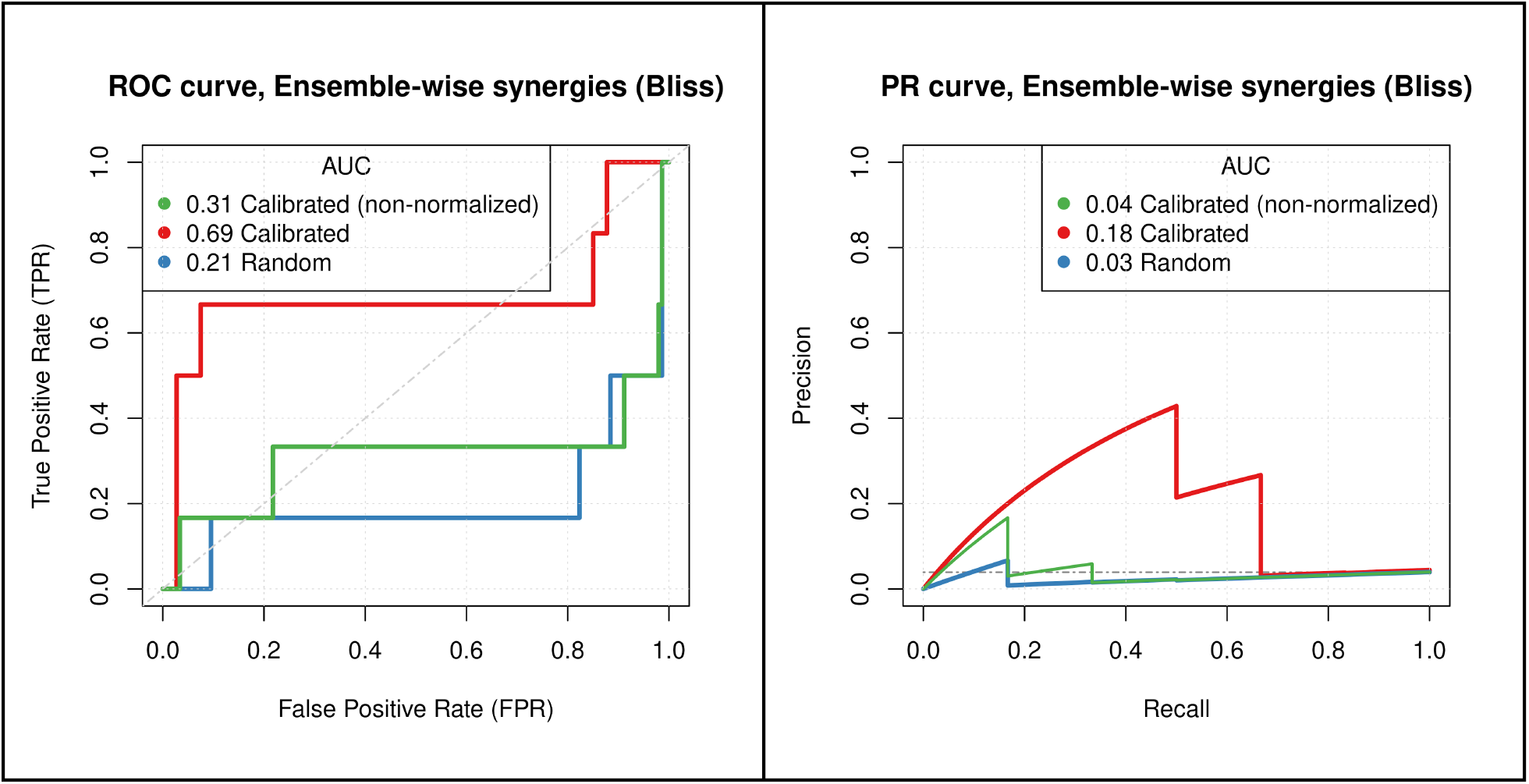
Predictive performance for random (proliferative) models, calibrated models (non-normalized) and calibrated normalized to random models (CASCADE 2.0 topology).

**Figure S3:**
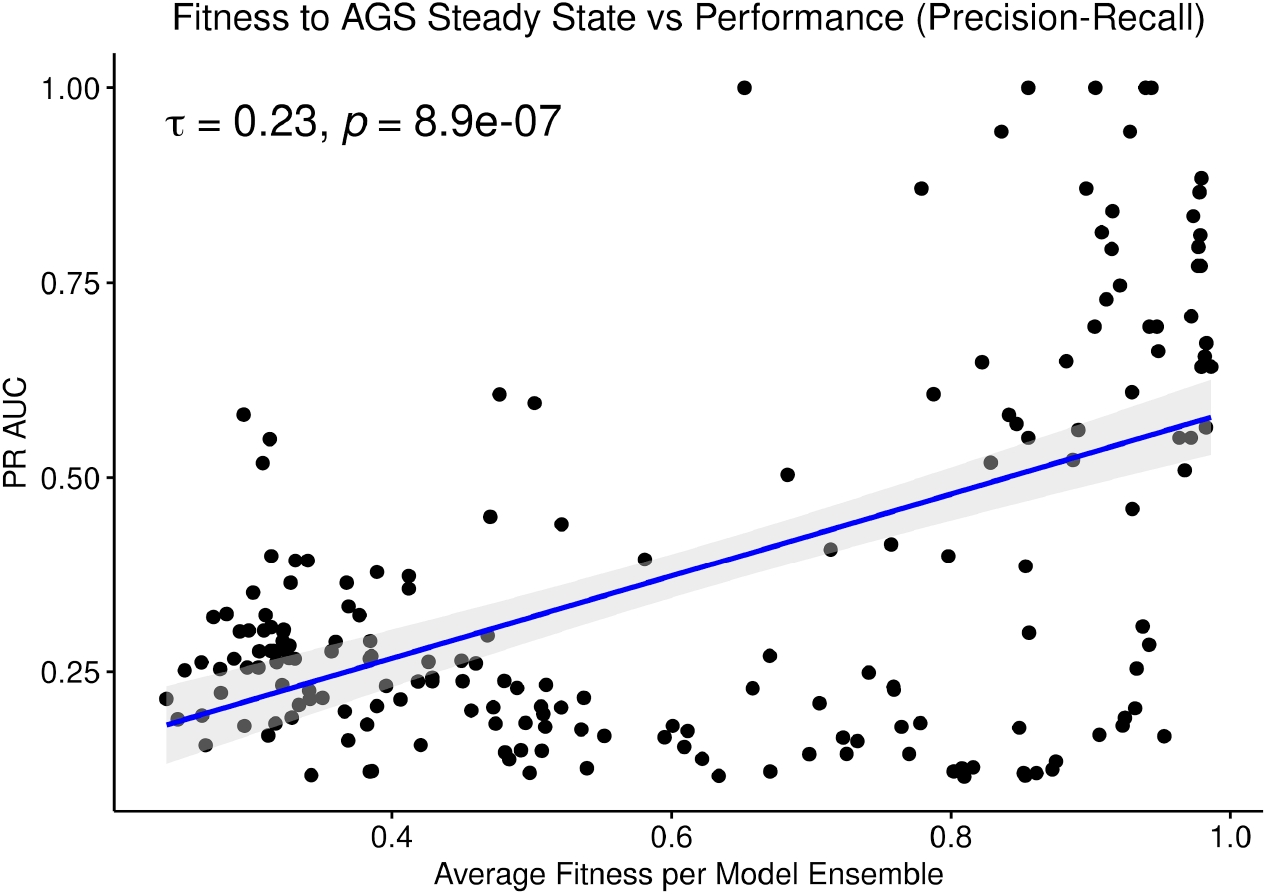
PR AUC performance dependence on fitness (CASCADE 1.0 topology).

**Figure S4:**
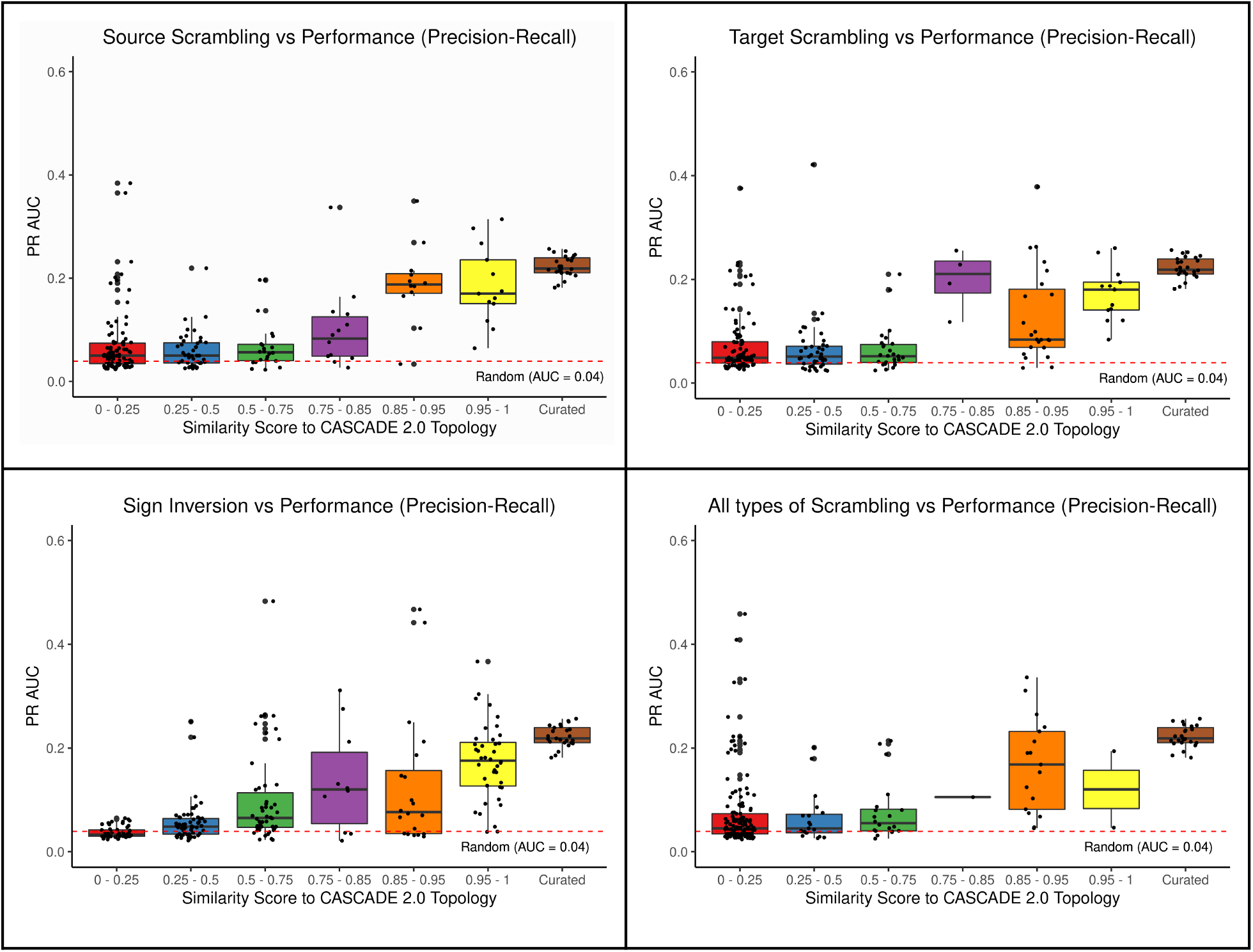
Effects of variations introduced in the CASCADE 2.0 prior knowledge graph (PR AUC performance metric).

**Figure S5:**
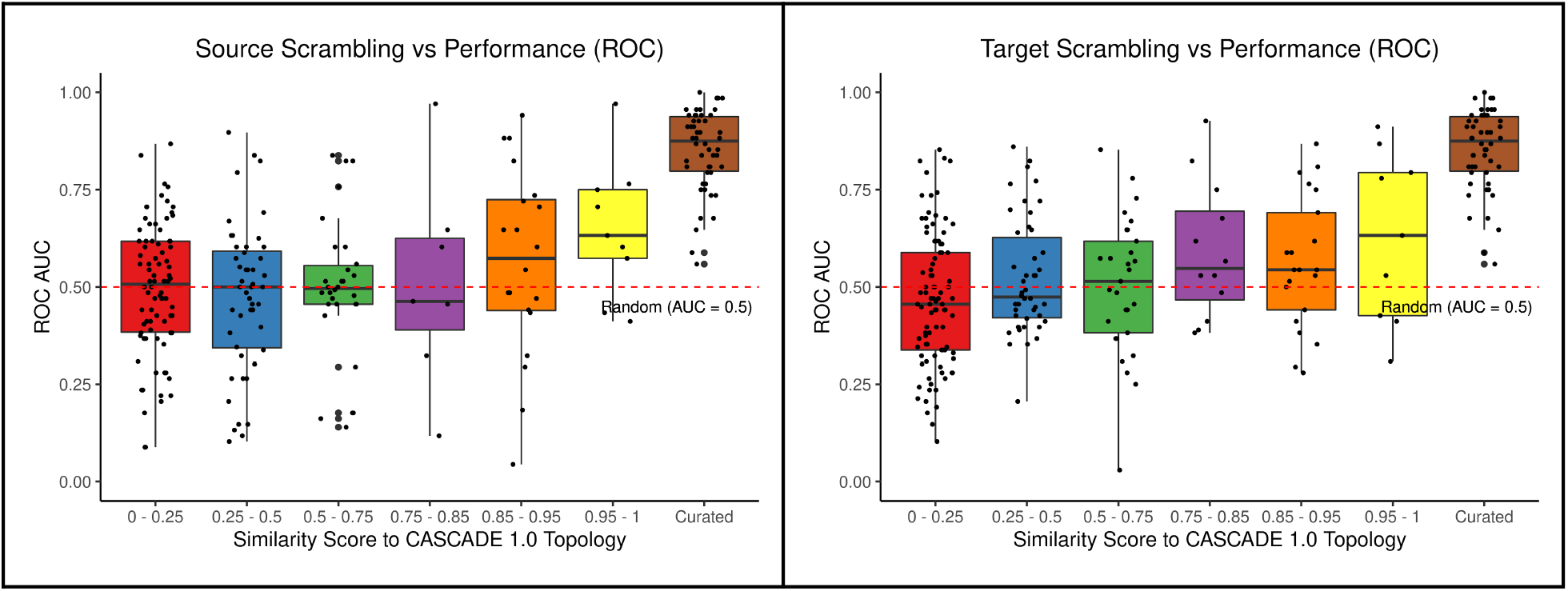
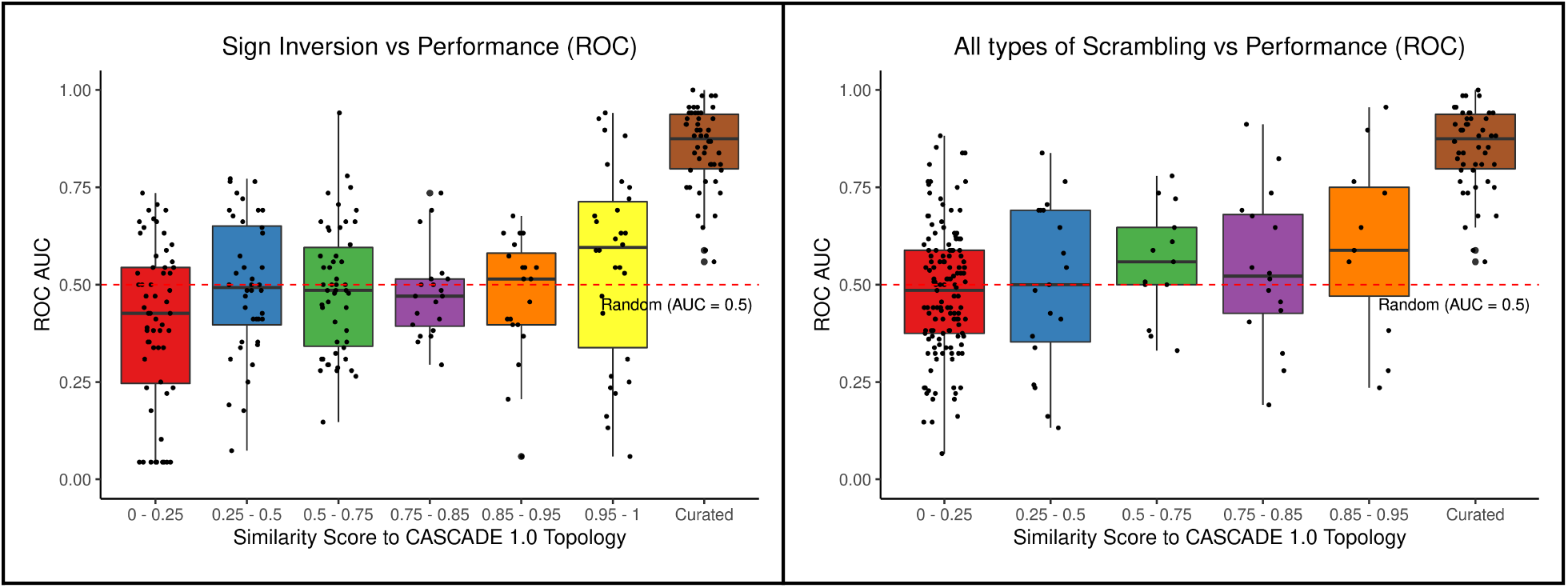
Effects of variations introduced in the CASCADE 1.0 prior knowledge graph (ROC AUC performance metric).

**Figure S6:**
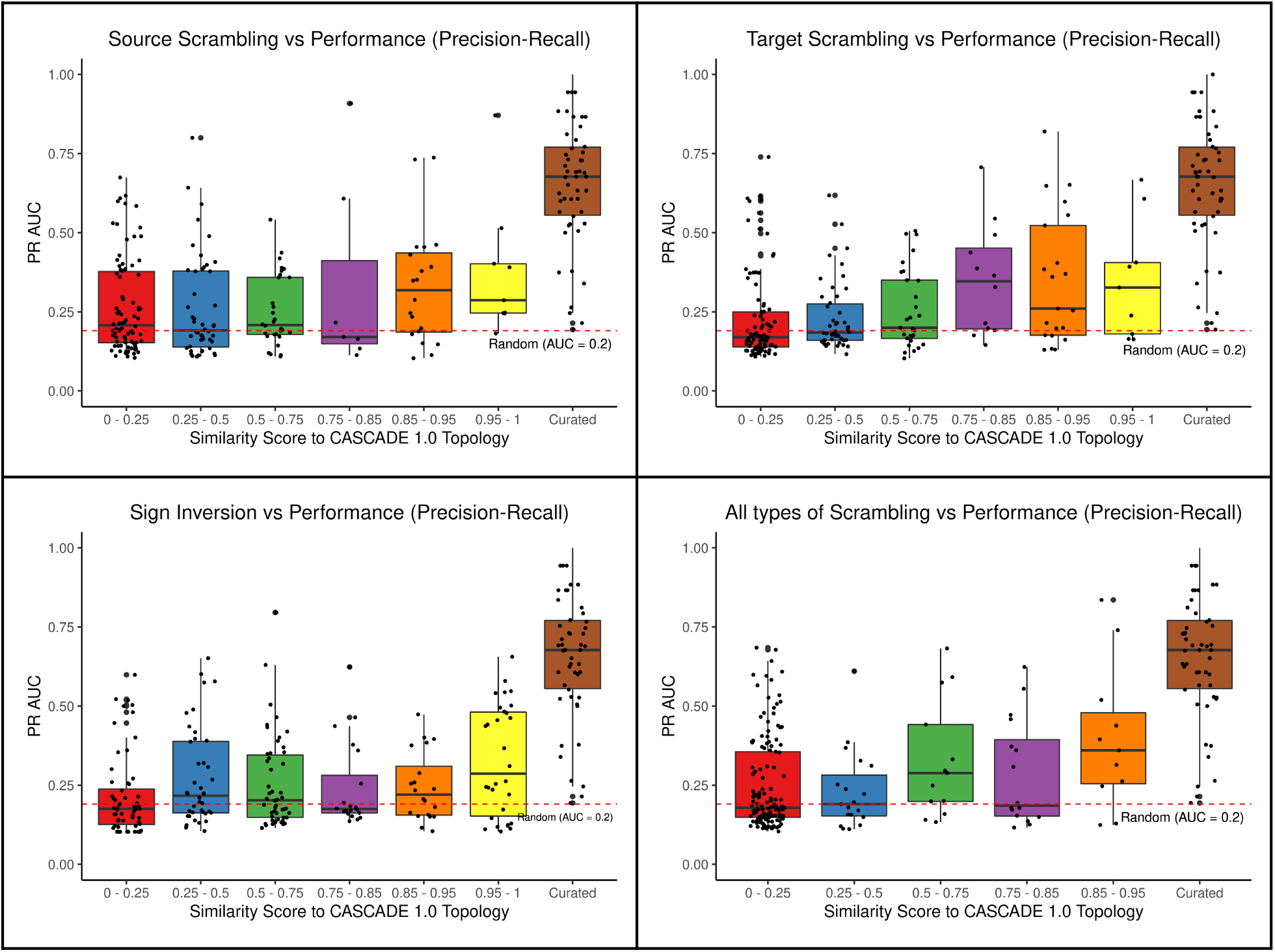
Effects of variations introduced in the CASCADE 1.0 prior knowledge graph (PR AUC performance metric).

